# Enhanced survival and low proliferation marks multifunctional virus-specific memory CD4 T cells

**DOI:** 10.1101/2022.12.13.520219

**Authors:** Lotus M Westerhof, Jonathan Noonan, Kerrie E Hargrave, Elizabeth T Chimbayo, Zhiling Cheng, Thomas Purnell, Mark R Jackson, Nicholas Borcherding, Megan KL MacLeod

## Abstract

Cytokine production by memory T cells is a key mechanism of T cell mediated protection. However, we have a limited understanding of the survival and secondary responses of memory T cells with cytokine producing capacities. We interrogate antigen-specific CD4 T cells using a mouse influenza A virus infection model. CD4 T cells with the capacity to produce cytokines survive better than non-cytokine+ cells, displaying a low fold contraction and expressing high levels of pro-survival molecules, CD127 and Bcl2. Transcriptomic analysis reveals a heterogenous population of memory CD4 T cells with three clusters of cytokine+ cells. These clusters match flow cytometry data revealing an enhanced survival signature in cells capable of producing multiple cytokines. These multifunctional cells are, however, less likely to proliferate during and following primary and secondary infections. Despite this, multifunctional memory T cells form a substantial fraction of the secondary memory pool, indicating that survival rather than proliferation may dictate which populations survive within the memory pool.

**Graphical Abstract.**
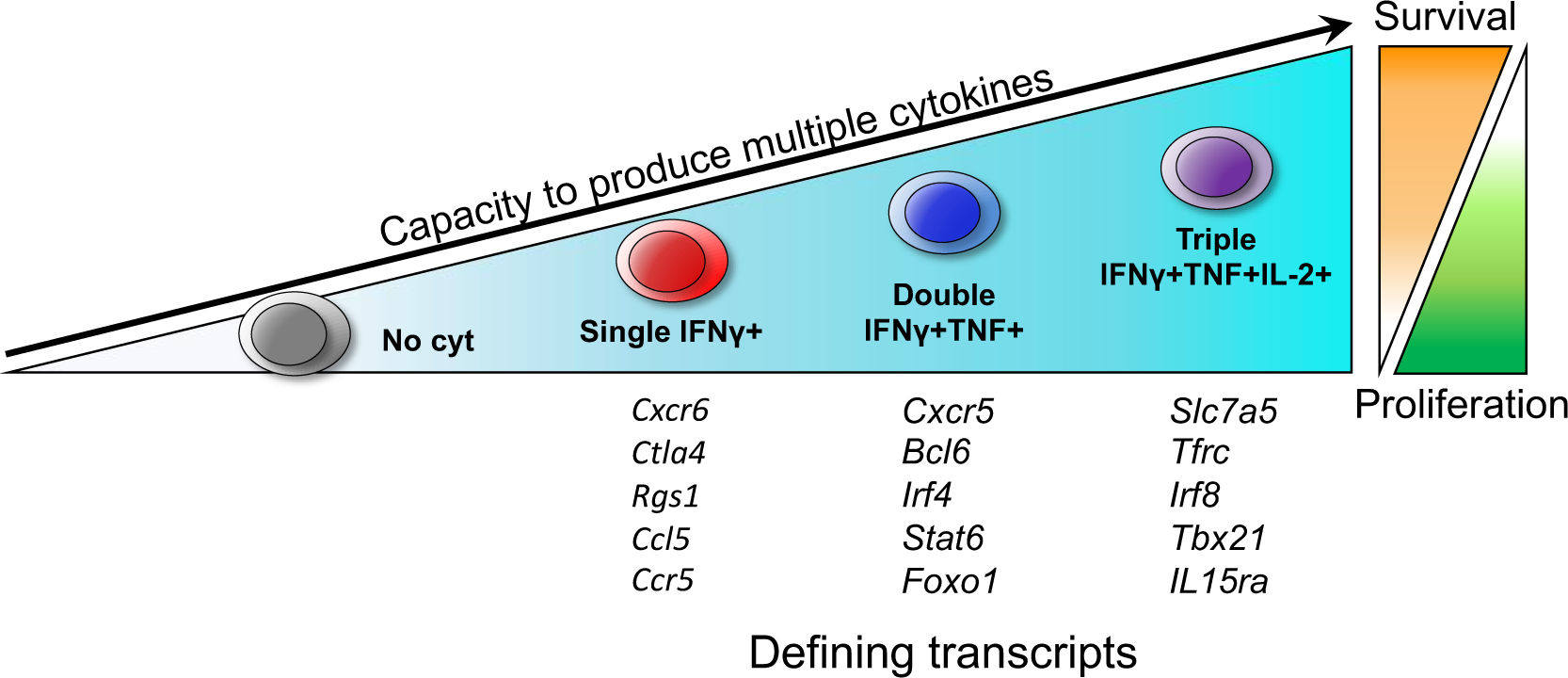

## Introduction

Immunological memory protects against repeated infection with a strong and rapid pathogen specific immune response. Cytokine production by memory T cells is often central to this protective response and the presence of T cells that can produce multiple different cytokines is associated with the most effective immune protection^1–10^. Understanding which memory T cells provide the most effective protection and the signals that generate these cells is a key target towards producing more effective vaccines.

Improving our understanding of T cell memory is particularly important in the context of highly variable infections such as Influenza A Virus (IAV). While neutralising antibodies can prevent IAV infection, the frequent mutations within viral surface proteins makes it difficult for these strain specific antibodies to recognise altered viruses^11^. As current IAV vaccines also induce antibodies that target strain specific surface proteins, they induce limited protection against different serotypes. This mechanism of escaping humoral immunity is also observed in other pathogens including HIV and SARS-CoV-2^12, 13^.

We know cytokine producing CD4 and CD8 T cells are key in protection against respiratory viruses, including IAV, at primary infection and upon reinfection with the same or altered viral strains. Importantly, CD4 and CD8 T cells recognise epitopes that are often conserved between IAV strains^14, 15^. The presence of cytokine producing IAV-specific CD4 and CD8 T cells in human peripheral blood correlates with cross-strain protection against symptomatic influenza infection^16–19^. Mechanistic mouse studies have further demonstrated the protective ability of IAV specific CD4 and CD8 T cells, with the inflammatory T-helper (Th1) cytokine, interferon-γ, often key to this response^2, 8, 9, 20–22^. The cytokines TNF and interleukin (IL-) 2 are also implicated in protection to IAV^23, 24^. These data support that the most effective memory T cells are those with the capacity to produce cytokines, in particularly multifunctional cells that can produce several different cytokines.

Memory T cells with the capacity to produce effector cytokines such as IFNγ and TNF can be classified as effector memory T (Tem) cells that can migrate through the blood and non-lymphoid tissues^25, 26^. In contrast, central memory T (Tcm) cells that migrate through secondary lymphoid organs can be capable of strong proliferation and are multipotent, differentiating into different functional subsets including effector cytokine producing cells and T follicular helper cells^27, 28^.

Cytokine producing memory T cells can also be found within populations of memory T cells located in non-lymphoid tissues that do not recirculate. These tissue resident memory T cells (Trm) can provide rapid protection to re-infection. These studies suggest vaccines should aim to induce Trm cells^29, 30^. However, heterosubtypic immunity to IAV, mainly mediated by CD4 and CD8 T cells, wanes in the months following infection of mice^21^. Most studies agree that lung memory CD8 and CD4 T cells decline over time, although recruitment of circulating cells or homeostatic proliferation may maintain these populations to some extent^31–35^. A long-lived protective memory response may, therefore, also be dependent on the generation of memory T cells beyond the pathogen-targeted tissue.

Fundamentally, a deeper understanding of protective memory T cells is required to reveal which memory T cells vaccines should aim to generate. We previously showed that multifunctional CD4 and CD8 T cells increase in predominance from the primary to the memory pool^36^. Building on this finding, we have now characterised the generation, survival, and function of these memory T cells following IAV infection in multiple tissues.

We define that a low fold contraction and a low level of proliferation by multifunctional T cells defines their increased predominance in the memory pool. Moreover, focusing on the CD4 compartment, we show T cells producing multiple cytokines demonstrate a strong pro-survival transcriptional signature. However, despite this pro-survival signature, multifunctional T cells proliferated poorly during a secondary IAV challenge *in vivo*, in contrast to single IFNγ+ T cells. By comparing memory CD4 T cells that do or do not have the capacity to produce cytokines, we reveal that cytokine producing T cells have an enhanced capacity to survive into the memory pool. Consequently, our study challenges the memory T cell paradigm that less differentiated central memory T cells have enhanced survival compared to more differentiated effector memory T cells^25, 26, 37, 38^.

## Results

### MHC tetramers and ex vivo cytokine analysis enable a comparison of CD4 and CD8 T cell phenotype and function following IAV infection

We used MHC I and MHC II tetramers (tet) containing immunodominant IAV nucleoprotein (NP) peptides, (NP_368-74_ and NP_311-325_) to identify IAV-specific T cells longitudinally following IAV infection, Figure 1A-B; gating shown in Supplementary Figure 1. These same peptides were used to separately activate cells from the same mice *ex vivo* to examine the anti-IAV T cell cytokine responses, Figure 1B, as optimal dual staining of MHC tetramers and cytokines was not possible. IAV infected animals were injected with fluorescently labelled anti-CD45 shortly before euthanasia to label cells in the blood; lung IAV-specific T cells that are CD45iv negative are likely Trm cells^39^.

**Figure 1:**
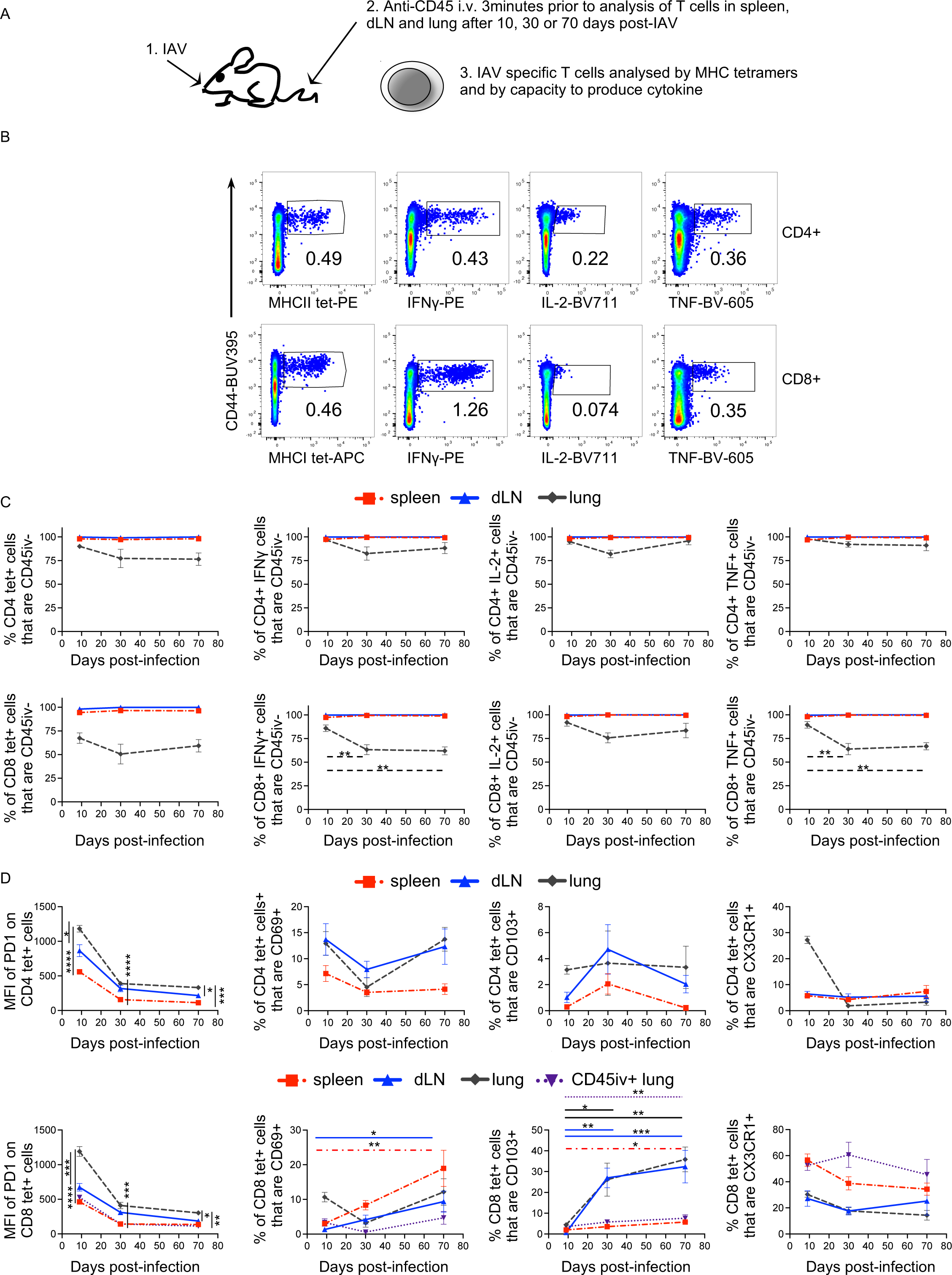
Memory IAV specific CD4 and CD8 T cells can be tracked using MHC tetramers and via ex vivo *cytokine responses*. C57BL/6 mice were infected i.n. with IAV on day 0 and injected i.v. with fluorescently labelled anti-CD45 3 minutes prior to removal of organs. Single cell suspensions of spleens, mediastinal draining lymph node (dLN), and lung were examined after 9, 30 or 70 days. Cells were either stained with IA^b^/NP_311-325_ or D^b^/NP_368-374_ tetramers or cells were restimulated in the presence of Golgi Plug for 6hours with NP_311-325_ and NP_368-374_ peptides loaded onto DCs (A). Example Flow cytometry plots of the IAV specific cells in the spleen, cells gated as shown in Supplementary Figure 1A (B). The percentages of IAV specific cells that bound to the i.v. injected anti-CD45 was analysed at each time point, cells gated as in Supplementary Figure 1B (C). The phenotype of MHC tetramer+ CD4 and CD8 T cells at each time point was examined in each organ (D). Data are from two independent time course experiments with a total of 7-8 mice/time point. In C-D, symbols show the mean of the group and error bars are SEM. Statistical difference tested by ANOVA followed by a Dunnett’s multiple comparison test for analyses between time points and by a Tukey’s test between organs, *: p<0.05, **:p<0.01, ***:p<0.001, ****:p<0.0001.

In the lymphoid organs, the majority of MHC tet+ and cytokine+ CD4 and CD8 T cells were negative for CD45iv, Figure 1C. The majority of lung IAV specific CD4 T cells were also not labelled with the i.v. injected anti-CD45 at any timepoints. In comparison, around 50-60% of the MHCI tet+ CD8 T cells were CD45iv+ at all time points and the proportion of labelled CD8 IAV specific IFN-γ+ and TNF+ T cells in the lung increased after day 9.

We examined the phenotype of the MHC tet+ T cells comparing cells in the spleen, dLN and lung across the three time points, Figure 1D and Supplementary Figure 1C-D. There were too few MHCII tet+ CD45+ lung CD4 T cells at memory time points to analyse reliably, but we have included data on CD45iv+ CD8 T cells.

As expected, PD1 was expressed on activated T cells and declined after day 9 on CD4 and CD8 T cells in T cells from all organs. However, MHC tet+ CD45iv negative CD4 and CD8 T cells in the lung continued to express higher levels than T cells from the lymphoid organs. The cell surface markers CD69 and CD103 are associated with Trm cells^29, 30, 40^. Expression of these molecules increased on CD8 MHCI tet+ cells in the lymphoid organs and CD45iv negative lung cells after day 9. However, very few CD4 MHCII tet+ cells expressed these markers.

While at day 9 some CD4 MHCII tet+ cells in the lung expressed CX3CR1, only a small minority of memory cells expressed this chemokine receptor. CX3CR1 was expressed by CD8 MHCI tet+ cells, most prominently in the spleen and on lung CD45iv+ cells suggesting that these are circulating effector memory cells^41^.

The phenotype of the lung CD45iv+ MHCI tet+ CD8 T cells more closely resembled the cells in the spleen rather than the CD45iv negative cells in the lung. The exception was in expression of CD69. While spleen MHCI tet+ CD8 T cells increased expression of CD69 across the time course, the circulating lung MHCI tet+ CD8 T cells remained low.

### T cells with the capacity to produce cytokines are more likely to enter the memory pool than non-cytokine+ T cells

We examined the decline of T cells detected with either MHC tetramers or IFNγ production following NP peptide restimulation. We focused on the decline between the peak response at day 9 to day 30, following the contraction phase^42^. As MHC tetramers detect NP specific T cells regardless of their ability to produce cytokines, altered dynamics of decline between these cells and those that are cytokine+ are suggestive of differences between cells that can and cannot produce cytokine.

CD4 MHC tet+ T cells declined in all three organs between day 9 and 30. In contrast, IFN γ+ cells did not decline in the secondary lymphoid organ, although these cells did decline in the lung (Figure 2A). Similarly, the numbers of IL-2+ and TNF+ did not decline in the spleen and dLN between day 9-30 (Supplementary Figure 2A). As for IFNγ+ cells, TNF+ cells declined in the lung but the numbers of IL-2+ cells were stable. These data suggest that, at least in secondary lymphoid organs, cytokine+ cells are less likely to be lost during the contraction phase than those that do not produce cytokine.

**Figure 2:**
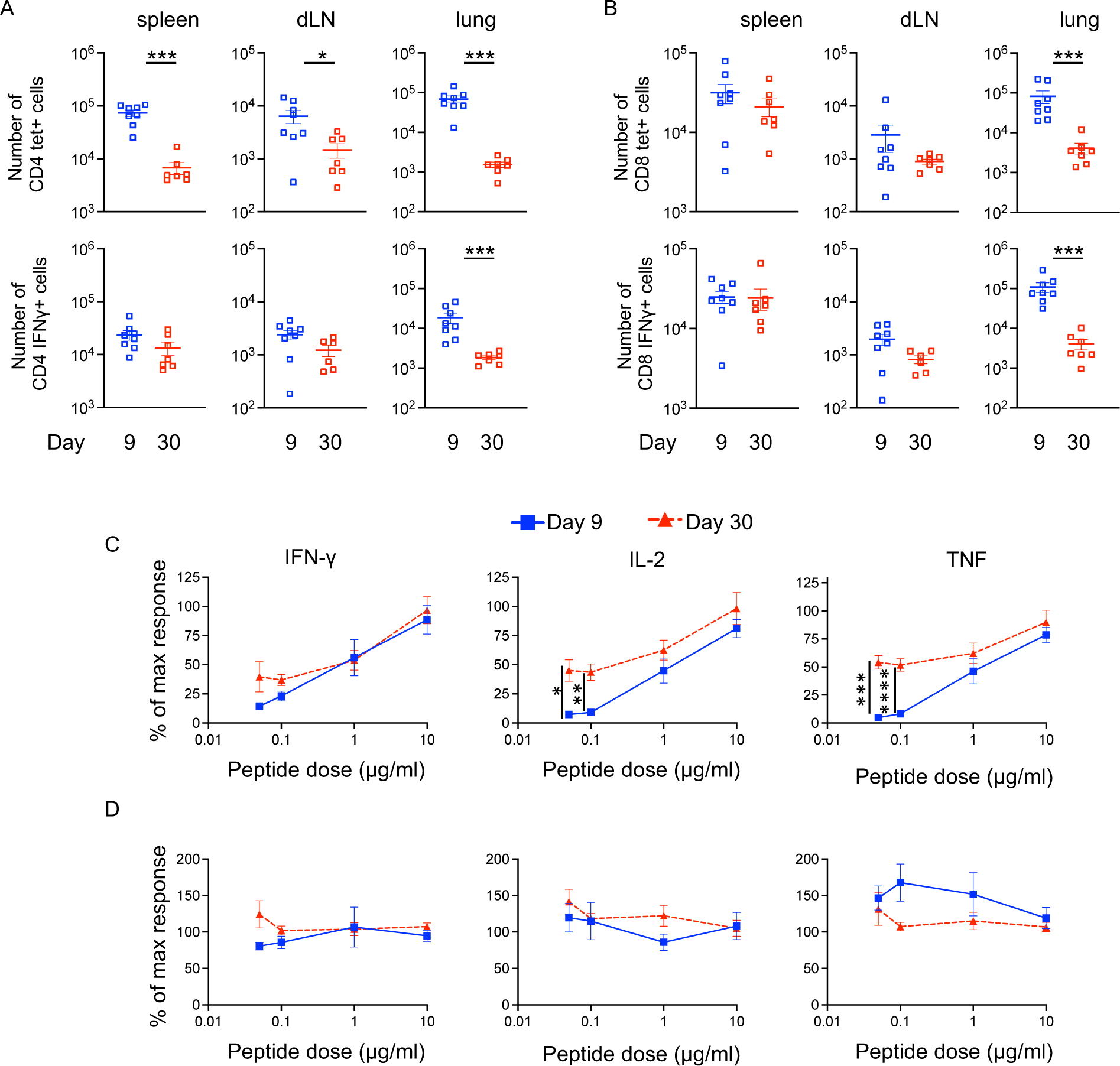
Influenza virus specific IFNγ+ CD4 T detected by MHC tetramers decline between day 9 and 30, while cytokine+ cells remain stable in secondary lymphoid organs. C57BL/6 mice were infected i.n. with IAV on day 0 and injected i.v. with fluorescently labelled anti-CD45 (CD45iv) 3 minutes prior to removal of organs. Single cell suspensions of spleens, mediastinal draining lymph node (dLN), and lung were examined after 9 or 30 days and either stained with MHCII/NP or MHCI/ NP tetramers, or activated with IAV-peptide loaded DC. Numbers of MHC tetramer or IFNγ+ CD4 (A) or CD8 T cells (B) were calculated. In C-D, spleen cells from C57BL/6 mice infected 9 or 30 days earlier with IAV were restimulated *ex vivo* with bone marrow DCs that had been previously incubated with 0.05, 0.1, 1, 10 or 100μg/ml of NP_311-325_ and NP_368-374_ for 2 hours. Co-cultures were for 6hours and contained GolgiPlug. Cells were gated on cytokine+ CD4 or CD8 T cells and the percentage of max cytokine production as detected at 100μg/ml calculated. In A-B, data are from two independent time course experiments with a total of 7-8 mice/time point. Y-axis set at the limit of detection and errors are SEM. Significance tested by a Mann-Whitney. In C-D, data are from two separate experiments with a total of 6-8 mice per timepoint. Significance between primary and memory tested by ANOVA followed by a Šídák’s multiple comparison test. In all data *: p<0.05, **:p<0.01, ***:p<0.001, ****:p<0.0001.

In contrast, IAV specific CD8 T cells detected by either MHC tetramers, IFNγ or TNF, remained stable between day 9 and 30 in secondary lymphoid organs, although in all cases these cells declined in the lung (Figure 2B and Supplementary Figure 2B). While the number of IL-2+ CD8 T cells were low, these cells were maintained at stable numbers in the spleen and lung, but did drop in the dLN.

Interestingly, for CD4 T cells, these changes were accompanied by an increase in sensitivity to peptide by the cytokine+ cells, with memory CD4 T cells producing TNF and IL-2 at lower peptide concentrations than cells at day 9. In contrast, primary and memory CD8 T cell cytokine production was unaffected by the peptide dose, Figure 2C-D. These data highlight important distinction between CD4 and CD8 T cells at primary and memory time points.

### Memory T cells with the capacity to produce cytokines express higher levels of pro-survival molecules than T cells that cannot produce cytokines

To examine why IAV specific CD4 cytokine+ and negative populations may display different dynamics we compared their phenotype and function by taking advantage of a reporter mouse we have developed^43^. In these triple transgenic, ‘TRACE’ mice, activated T cells express rtTA, which, when bound to doxycycline, activates the tet-ON promoter, driving Cre expression which leads to permanent EYFP production at the ROSA locus. Infection with IAV induced a substantial population of EYFP+ CD4 T cells when triple transgenic mice were fed a doxycycline+ diet during the initial infection, Supplementary Figure 3. This model greatly increases the number of responding T cells we can identify and enables identification of cytokine+ and negative EYFP+ cells within the same sample, Figure 3A-B.

**Figure 3:**
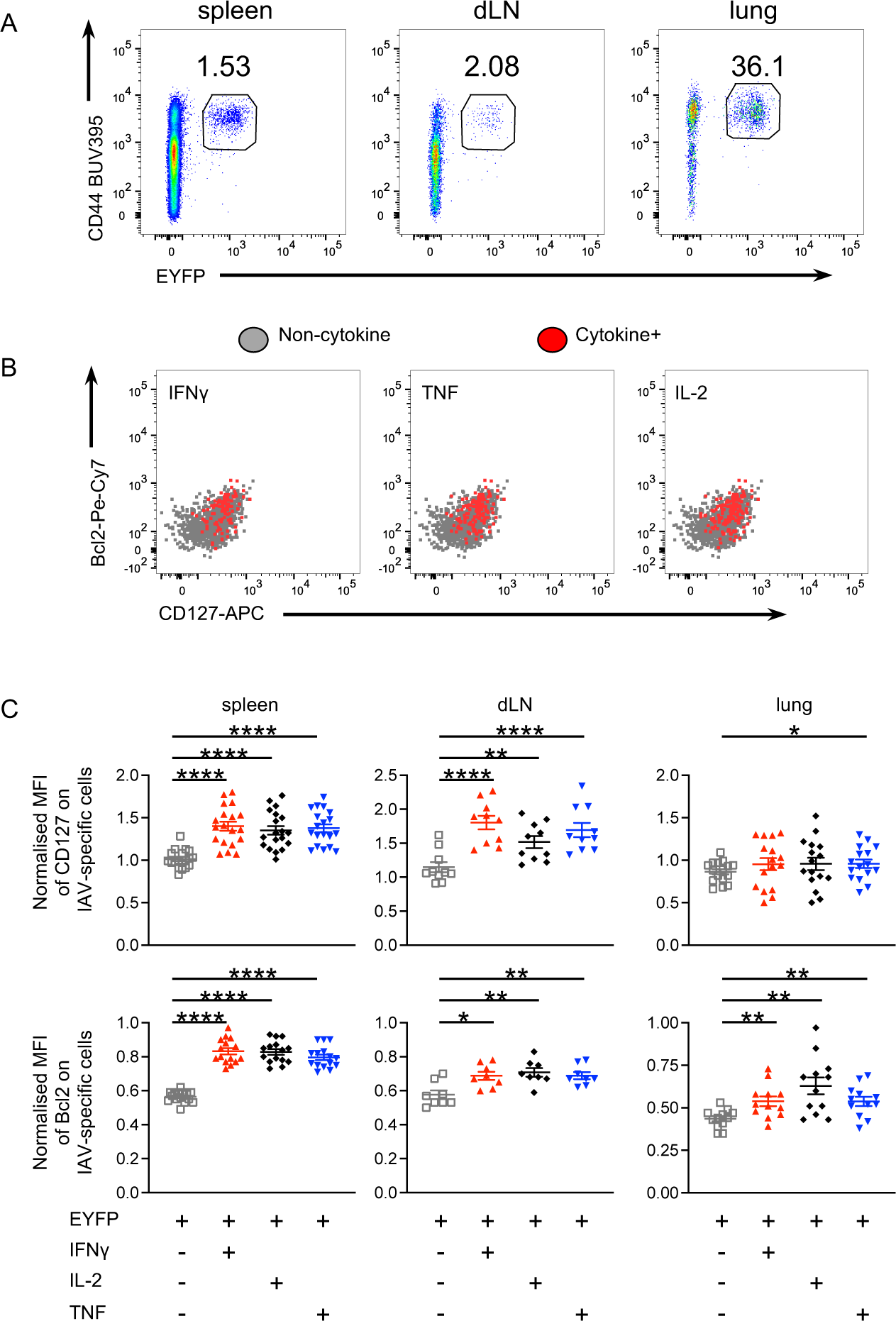
Cytokine+ memory CD4 T cells express higher levels of CD127 and Bcl2 than cytokine negative cells. TRACE mice were infected i.n. with IAV on day 0 and injected i.v. with fluorescently labelled anti-CD45 (CD45iv) 3 minutes prior to removal of organs at day 40. Single cell suspensions were re-activated with IAV Ag-loaded DCs. Example FACS plots from the indicated organs (A) and the spleen (B) are gated on live CD4+ dump negative, CD45i.v. negative cells (A) and on EYFP+ cytokine negative: grey; IFNγ, IL-2 and/or TNF+: red (B). In C, data are from three independent experiments with a total of 5-8 mice and normalised by dividing the MFI on EYFP+ cells by the MFI of naïve CD44^lo^ CD4 T cells from cells from the same mouse and organ. Symbols represent a mouse and the lines shows the means, error bars are SEM. Samples from some dLNs and lungs were excluded as the numbers of EYFP+ cells collected were too low for analysis. Significance tested via a Friedman paired analysis with Dunn’s multiple comparison test, *: p<0.05, **:p<0.01, ****:p<0.0001.

As the EYFP+ CD4 T cells are likely to respond to an array of IAV antigens, we used bone marrow derived dendritic cells (DCs) cultured with a sonicated IAV antigen preparation to restimulate the T cells *ex vivo* for cytokine analysis^36^ Supplementary Figure 3D. The expression of the pro-survival molecules CD127 and Bcl2 was examined in EYFP+ cytokine negative CD4 T cells and those expressing IFNγ, IL-2 and TNF following *ex vivo* restimulation of cells taken from mice infected 40 days previously, Figure 3B-C. In the secondary lymphoid organs, the cytokine+ EYFP+ T cells expressed higher levels of both molecules compared to cytokine negative cells. These differences were less clear in the lung, potentially reflecting the reduced survival of cytokine+ and negative cells in this organ, but all cytokine+ cells expressed significantly higher levels of Bcl2 than the non-cytokine+ cells. These data are consistent with the hypothesis that CD4 T cells with the capacity to produce cytokines have an enhanced survival capacity compared to non-cytokine+ T cells.

### T cells that can produce multiple cytokines are more persistent than those that only have the capacity to produce IFNγ

To investigate the cytokine+ T cells in more detail, we separated these into single IFNγ+, double IFNγ+ TNF+ or IL-2+ cells, or triple cytokine+ cells detected following reactivation with IAV Ag-loaded DCs. Triple cytokine+ CD4 T cells did not decline in any of the organs between days 9 and 30, Figure 4A. All other populations declined significantly in at least one of the organs examined although cells in the lymph node had limited decline regardless of which population was examined.

**Figure 4:**
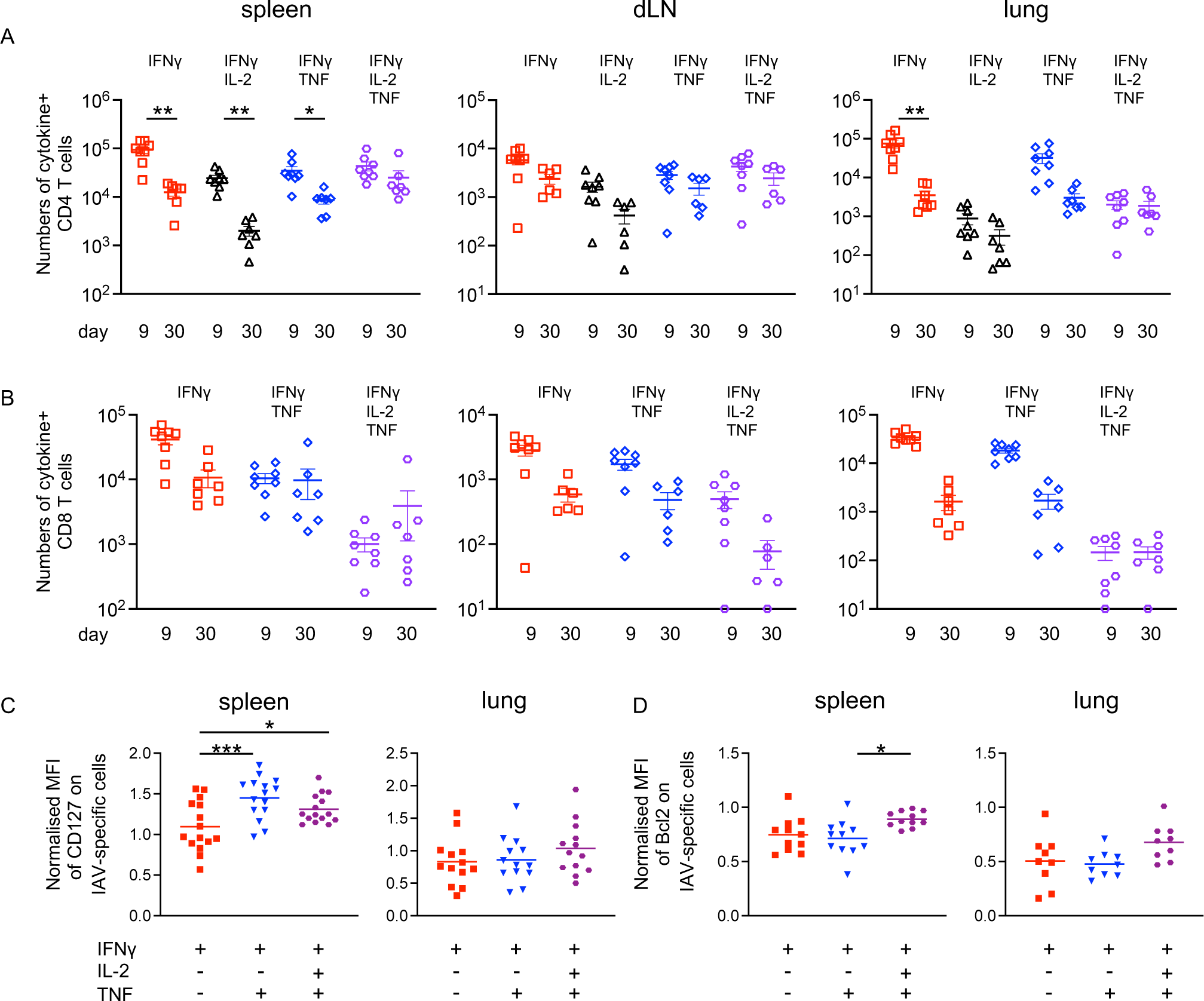
Triple cytokine+ cells are less likely to decline than single IFNγ+ T cells. C57BL/6 mice were infected i.n. with IAV on day 0 and injected i.v. with fluorescently labelled anti-CD45 3 minutes prior to removal of organs. Single cell suspensions of spleens, mediastinal draining lymph node (dLN), and lung were activated after 9 or 30 days post-infection by DCs incubated with IAV-Ag preparation. CD45iv negative IAV specific cytokine+ CD4 (A) and CD8 (B) T cells were detected by flow cytometry. In C-D, CD127 and Bcl2 expression on CD45iv negative cytokine+ CD4 T cells were detected by flow cytometry and normalised by dividing the MFI on the cytokine+ cells by the MFI of naïve CD44^lo^ CD4 T cells from the same mouse and organ. In A-B, data are from two independent time course experiments with a total of 7-8 mice/time point and the Y-axis is set at the limit of detection. Samples from some dLNs and lungs were excluded as the numbers of total cells collected were too low for analysis. Symbols represent each mouse, the bar shows the mean of the group and errors are SEM. Significance tested by a Kruskal-Wallis test followed by a Dunn’s multiple comparison test with Benjamini-Hockberg correction for multiple comparisons *: p<0.05, **:p<0.01, ***:p<0.001, ****:p<0.0001. In C-D, data are combined from 2-3 experiments with 4-5 mice per experiment, each symbol represents a mouse and the line shows the mean of the group, error bars are SEM. Significance tested via ANOVA with a Dunn’s multiple comparison test, *: p<0.05, ***:p<0.001, ****:p<0.0001.

There were no significant declines in the numbers of cytokine+ CD8 T cells, Figure 4B. However, in the lung, while triple cytokine+ CD8 T cells were clearly stable, the single IFN γ+ and double IFNγ+TNF+ cells displayed non-significant drops.

The triple cytokine+ T cells may display greater persistence than the single IFNγ+ T cells because they express higher levels of survival molecules. To address this, we compared the expression of pro-survival molecules, CD127 and Bcl2^44^, on single, double and triple cytokine+ cells. There were too few triple+ CD8 T cells in any organ and too few cytokine+ cells in the dLN to compare the MFI of the different populations. Therefore, we focussed on cytokine producing CD4 T cells in the spleen and lung.

At day 30 post-infection, we found that double and triple cytokine+ CD4 T cells in the spleen had higher expression of CD127 and triple+ cells in the spleen expressed more Bcl2 than double IFNγ+TNF+ cells, Figure 4C-D. While there was a trend for the triple cytokine+ cells in the lung to express higher levels of CD127 and Bcl2, this did not reach significance. These data suggest that the differences in these two pro-survival molecules may not be sufficient to explain the altered survival in lung cytokine+ memory T cells.

### Single IFNγ+ CD4 and CD8 T cells are more likely to be in cell cycle than multifunctional T cells

An alternative explanation for the enhanced stability of triple cytokine+ T cells was increased proliferation by the triple cytokine+ as compared to single IFNγ+ cells. We addressed this by examining Ki67, expressed during the cell cycle. We focussed on T cells detected following *ex vivo* reactivation with DCs presenting IAV antigens as these provided more robust populations than following peptide-restimulation. The numbers of IFNγ/IL-2 double+ T cells were low and therefore were not included in the analysis.

In the dLN at day 9, over 20% of the CD4 and 30% of the CD8 cytokine+ T cells were Ki67+. In contrast, in the spleen and lung, less than 15% of the CD4 and 25% or less of CD8 cytokine+ cells were Ki67+, Figure 5A-B. As expected, the percentages of T cells that were Ki67+ decreased from day 9 to day 30 and very few Ki67+ cells could be detected on day 70.

**Figure 5:**
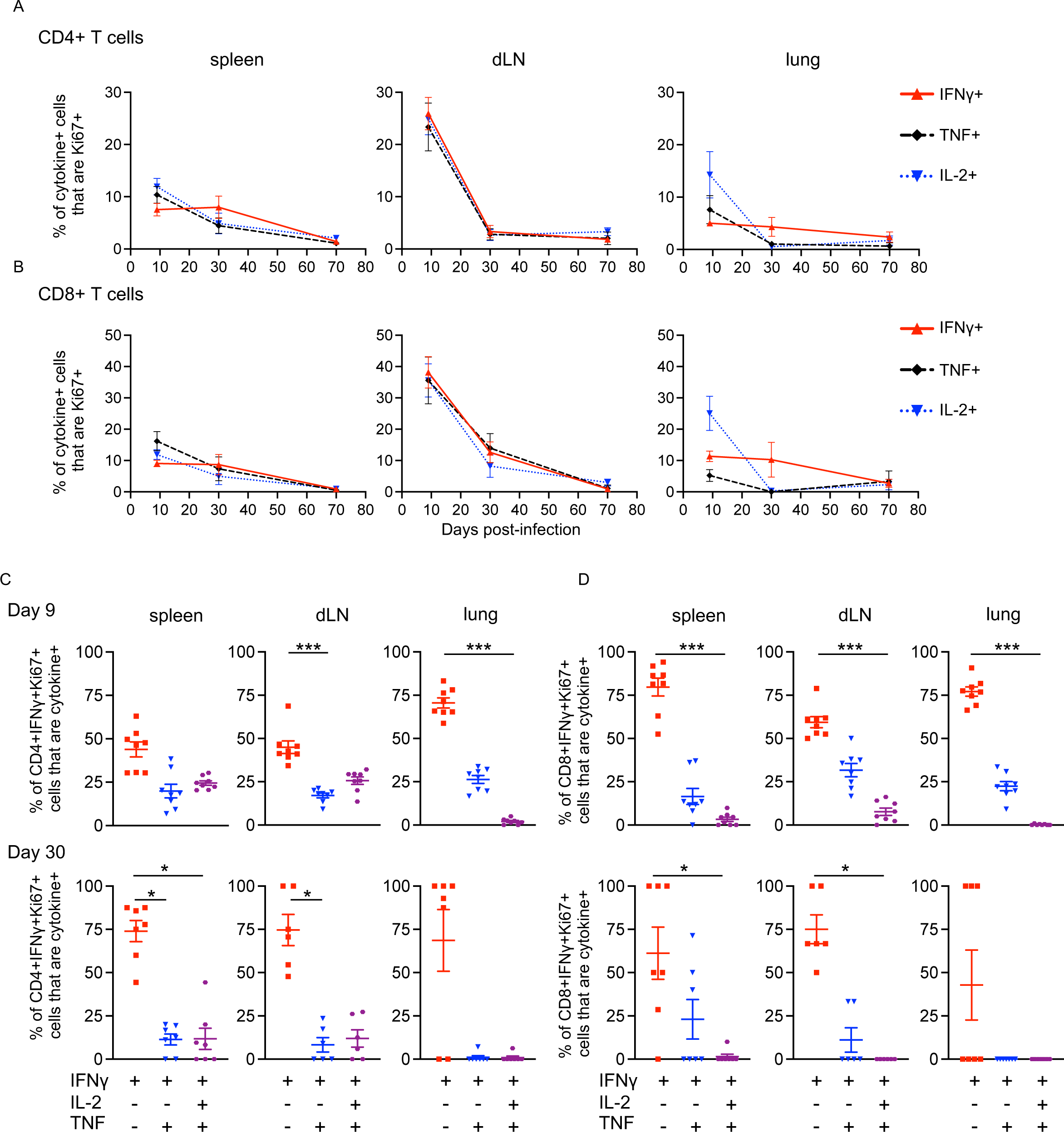
Single IFNγ+ CD4 and CD8 T cells are more likely to be in cell cycle 9 and 30 days after IAV infection than T cells producing multiple cytokines. C57BL/6 mice were infected i.n. with IAV on day 0 and injected i.v. with fluorescently labelled anti-CD45 3 minutes prior to removal of organs at the indicated time point. Single cell suspensions of spleens, mediastinal draining lymph node (dLN), and lung were activated by DCs incubated with IAV-Ag preparation. CD45iv negative cytokine+ T cells were detected by flow cytometry to detect Ki67 expression by cytokine producing T cells (CD4: A, C; CD8: B, D). Data are from two independent time course experiments with a total of 7-8 mice/time point. In C-F, each symbol represents a mouse and the horizontal line shows the mean of the group and significance tested via paired Friedman analysis with Dunn’s multiple comparison test *: p<0.05, ***:p<0.001.

We gated on the IFNγ+ Ki67+ cells to examine whether these cells were producing one or more of the cytokines. We hypothesised that triple cytokine+ cells may be more likely to proliferate to maintain their numbers in the memory pool. In contrast, we found the opposite result: IFNγ single+ CD4 and CD8 T cells were more likely to be in cell cycle compared to triple+ cells at both day 9 and 30, Figure 5C-D. These data rule out the hypothesis that triple cytokine+ cells increase proportionally in the memory pool compared to the primary pool because they proliferate more than single IFNγ+ cells.

### Single cell RNAseq analysis reveals a heterogeneity in cytokine+ and cytokine negative CD4 memory T cell populations

To obtain a more detailed understanding of differences between the cytokine positive and negative memory T cell populations, we performed single-cell gene transcription analysis. CD4 T cells from the spleens and lungs from TRACE mice infected with IAV 40 days previously were activated with IAV-Ag DCs for 4 hours to reveal those with the capacity to produce cytokines. CD45iv negative, EYFP+ CD44^hi^ cells from the spleen and lung and control naïve (CD44^lo^/EYFP negative) spleen cells were FACS sorted and gene expression examined via BD-Rhapsody single-cell transcriptomic analysis using the Immune Response Targeted panel for mouse.

Initial clustering produced 19 different clusters, including 8 naïve/central memory clusters, 10 memory clusters and one Treg cluster, Supplementary Figure 4A. To focus on the differences between the memory clusters, we reduced the complexity of the naïve/CM population into a single population, Figure 6A; all clusters remained transcriptionally distinct, Supplemental Figure 4B; Supplementary Table 1. Of the 10 memory clusters, 7 expressed little to no *Ifng*, *Tnf or Il2*, Supplemental Figure 4C. Cells from all clusters were found in both the spleen and lung, with the spleen contributing most cells in all but one cluster. The exception was the single+ cluster expressing *Ifng* but not *Tnf* or *Il2*, which was predominantly derived from the lung, Figure 6A & Supplementary Figure 4D.

**Figure 6:**
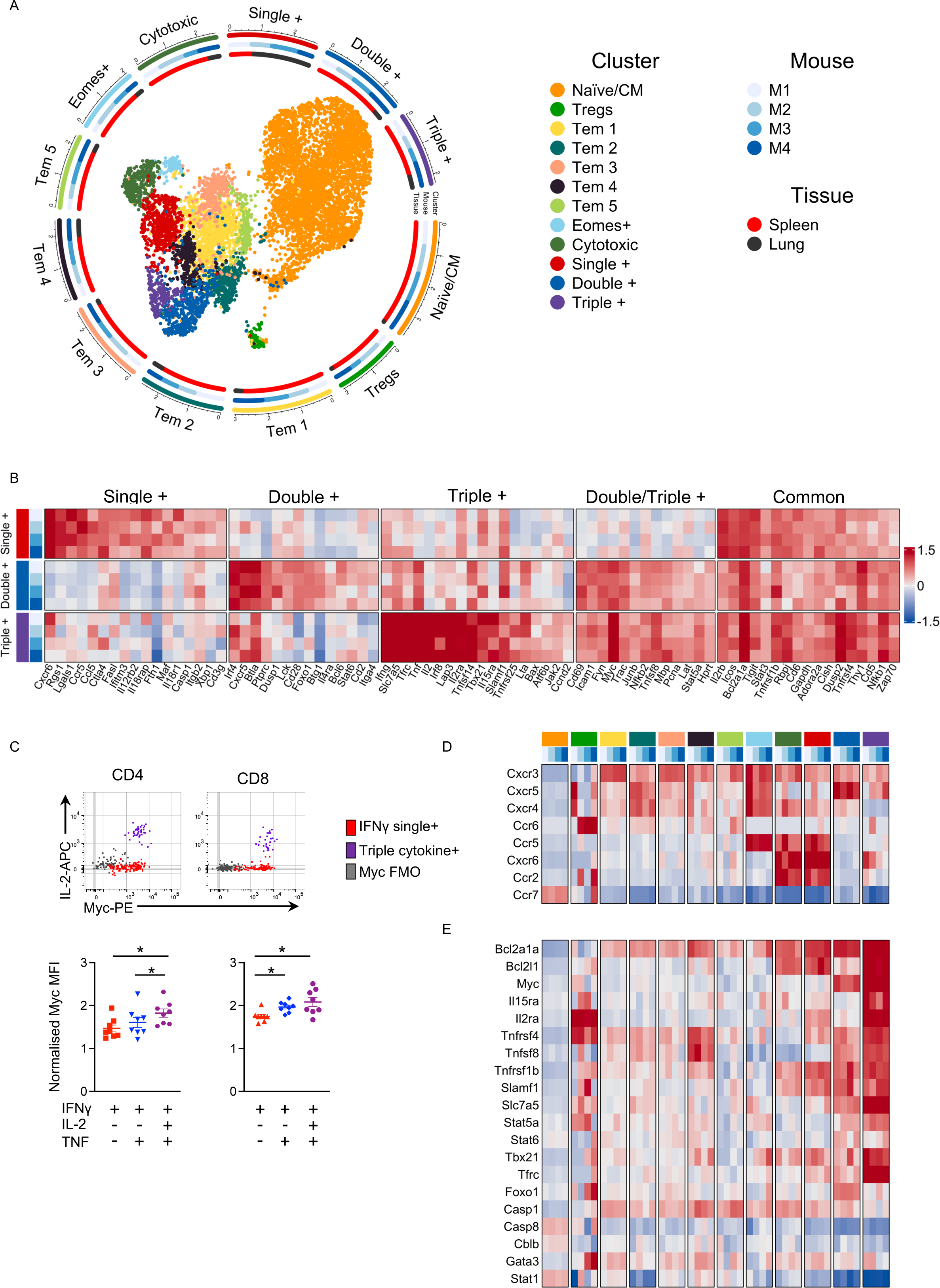
Triple cytokine+ CD4 T cells have a pro-survival transcriptional signature. TRACE mice were infected with IAV and 40 days later injected with anti-CD45 3minutes prior to removal of spleens and lungs. Isolated CD4 T cells were activated for 4 hours with IAV-Ag DCs and the CD45iv negative CD4+CD44^hi^EYFP+ cells (spleens and lung) and CD4+CD44^lo^EYFPnegative (spleens) cells were FACS sorted and their transcriptomes examined by scRNAseq. UMAP of one naïve/central memory cluster, one Treg cluster, and 10 memory clusters. The radial tracks around the UMAP reflect the numerical representation of each cluster, and the mouse/tissue origin within each cluster on a logarithmic axis (A). DEGs in single, double, and triple cytokine+ clusters identified by comparison between the indicated populations and all other clusters (B). The normalised MFI of Myc by single, double, and triple cytokine+ cells in CD4 and CD8 T cells from C57BL/6 IAV infected mice examined at day 40 post infection; expression was normalised by dividing each sample’s MFI by the MFI on naïve CD4+ cells in same sample (C). Expression of mRNA for chemokine receptors and with cell survival molecules from the scRNAseq analysis (D,E). Data in A, B, D, and E are from one experiment with cells from 4 mice. In C, data are from 2 independent experiments with each symbol representing one mouse and the horizontal line showing the mean of the groups; significance tested by paired ANOVA with Tukey’s multiple comparison test, *: p<0.05.

Triple+ cells were uniquely enriched for *Il2* and, while the single+ and double+ clusters clearly expressed *Ifng* +/− *Tnf*, these transcripts were only significantly enriched (versus all other cells) within the triple+ cluster. Previous studies have demonstrated that multifunctional T cells produce more IFNγ on a per cell basis than single IFNγ+ T cells^1, 3, 6^. We also found that triple cytokine+ T cells express the greatest amounts of IFNγ by flow cytometry, confirming our scRNAseq data, Supplementary Figure 5. Moreover, by gating on total IL-2+ or TNF+ T cells, we found that triple cytokine+ CD4 T cells also expressed more of these cytokines than single IL-2+ or TNF+ cells respectively. Interestingly double cytokine+ CD4 T cells mainly displayed an intermediate phenotype with a tendency to produce more of the cytokine than single producers but less than triple+ T cells.

### Triple cytokine+ memory CD4 T cells express a pro-survival gene signature

We have focussed on the transcriptional signatures of the three cytokine producing clusters: single+ (*Ifng*) double+ (*Ifng* & *Tnf*), and triple+ (*Ifng, Tnf* & *Il2*) cells, Figure 6B & Supplementary Table 1. The single+ CD4 T cells expressed the highest levels of *Icos* and *Pdcd1* (PD1), which we confirmed at protein level by flow cytometry, Supplementary Figure 6. These cells also expressed other transcripts implicated in T cell activation including *Ctla4*, *Tnfrsf9* (41BB), and *Ccl5*. High expression of *Cxcr6* and *Rgs1* may reflect that many of these were isolated from the lung^45^ and expression of *Xbp1* tallies with the proliferative ability of these cells^46^.

Genes uniquely upregulated in the double+ cells indicated a Tfh-like phenotype, supported by the expression of *Cxcr5*, *Bcl6, Irf4*^47, 48^ and *Foxo1*^49^, suggesting these cells may be particularly important in supporting the antibody response in either the spleen or lung^50, 51^. In addition to the increased transcription of cytokine genes, triple+ cells were also enriched for genes such as*Tnfsf14* (LIGHT), *Tbx21* (T-bet), *Il2ra* and *Irf8*.

We also observed genes commonly enriched across some or all cytokine+ clusters. For example, the double and triple cytokine+ cells had overlapping expression of several genes reflective of T cell activation *Cd69*, *Icam1*, *Fyn* and *Stat5a*. Strikingly, the double and triple cytokine+ cells expressed high levels of the transcription factor *Myc* and we confirmed increased Myc protein expression in triple+ cells by flow cytometry, Figure 6C.

*Myc* is a key regulator of T cell metabolism downstream of activation and may confer both proliferative and/or survival advantages to these cells^52^. In addition to being the highest expressors of *Myc,* triple+ cells were also enriched for *Myc* pathway associated transcripts, expressing the highest levels of *Ybx3*^52^ and being uniquely enriched for *Tfrc* (CD71) and *Slc7a5* within the cytokine positive populations. Collectively, these data suggest that these T cells have a distinct metabolic profile compared to the other memory populations.

Interestingly, the different clusters of memory T cells expressed varying combinations of chemokine receptors suggesting that they may be located in distinct regions of the lung and spleen and/or respond distinctly following re-infection, Figure 6D. Finally, the triple cytokine+ memory CD4 T cells expressed high levels of molecules associated with survival, Figure 6E. These included Bcl2 family members (*Bcl2a1a*^53, 54^ and *Bcl2l1*(Bclx)^55^), receptors for pro-survival cytokines (*Il2ra*, *Il15ra*^56^) and costimulatory molecules (*Tnfrsf4* (OX40), *Tnfsf8* (CD30L) associated with T cell survival^57–59^. Surprisingly, the cytokine+ cells expressed low levels of *Il7r* (CD127) which is in contrast to our flow cytometry data, Supplementary Figure 4B. This may reflect that gene transcription does not always relate to protein expression^60^.

### TCR clones are found in multiple memory T cell clusters but are not shared between different animals

We used CDR3 sequencing of the T cells’ TCRs to address whether clones were restricted to particular memory T cell clusters within the scRNAseq data. We found expanded clones within each of the memory populations, with the relative absence of these in the naïve/CM and regulatory populations, Figure 7A-B. Few expanded clones appeared cluster specific, with most represented across multiple clusters, Figure 7C-D. These data suggest that TCR CDR3 sequence does not dictate T cell fate. Finally, analysis of expanded clonotypes derived from each animal demonstrated little evidence of public T cell clones, Figure 7E.

**Figure 7:**
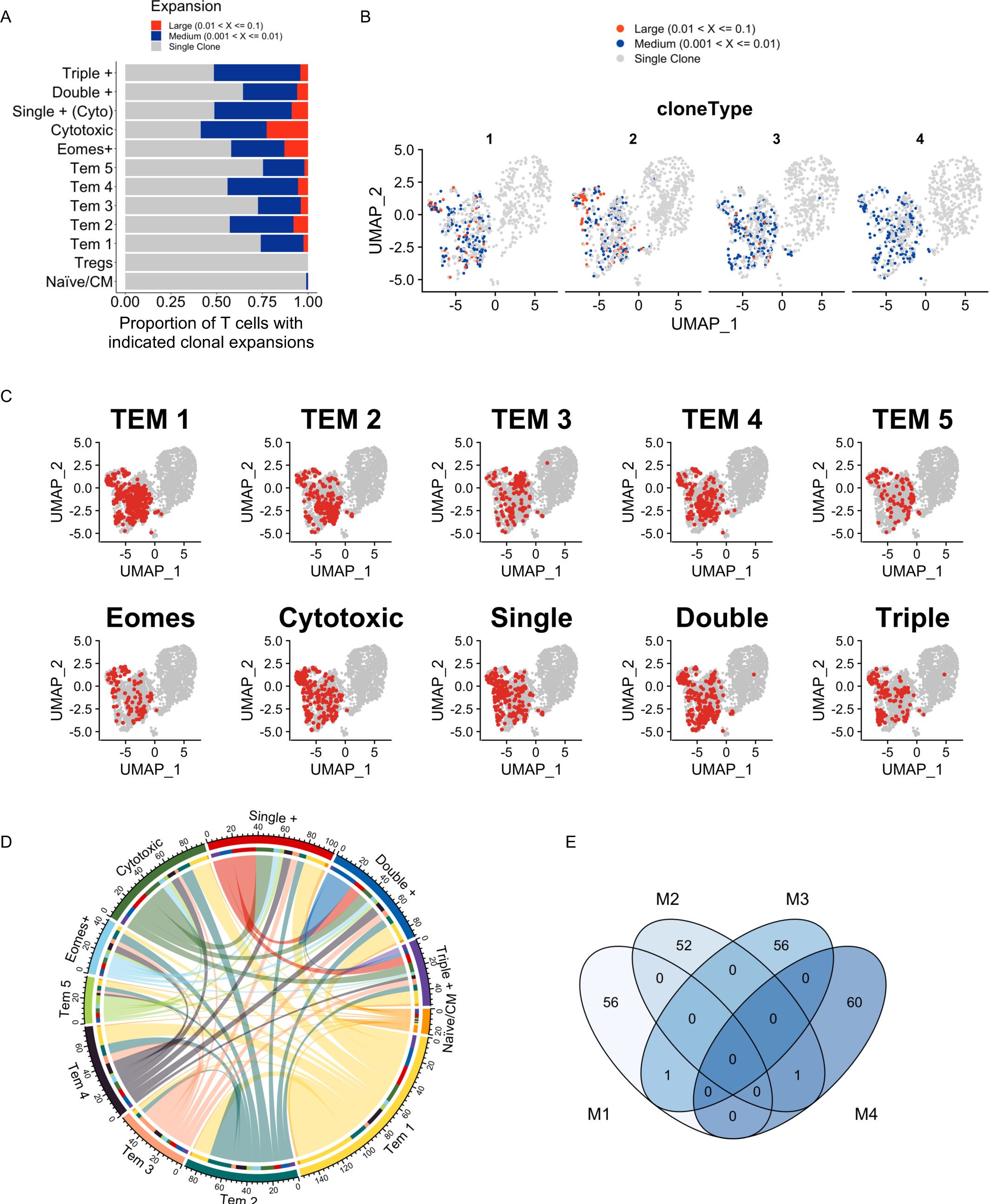
TCR clones are found in multiple memory T cell clusters but are not shared between different animals. scRNAseq analysis described in Figure 5 was integrated with single cell TCR CDR3 analysis. Large, medium and single clones identified by TCR CDR3 sequencing are displayed for each cluster and the proportion of T cells with the indicated clonal expansions shown (A). Enriched clones are overlaid on the scRNAseq UMAP for each animal and coloured by the magnitude of expansion (B). To highlight the extent to which clones were shared across clusters, clones found within each memory cluster were identified and displayed on the UMAP (C). To provide higher resolution these data are presented as a chord diagram, highlighting the inter-cluster relationship of individual clones (D). The limited overlap of identified clones between mice is displayed as a Venn diagram (E).

### Single IFNγ+ T cells dominate the proliferative response during re-infection

Our data above and the relationship between T cell multifunctionality, persistence into the memory pool, and protection from disease, suggest that triple cytokine+ T cells should dominate a secondary response to IAV^1, 3–6, 36^. In particular, the high expression of Myc by the triple and double cytokine+ cells suggested these cells would respond and proliferate robustly following reactivation^52^.

To test this, we re-infected IAV-memory mice with a distinct strain of IAV, X31, which has different surface proteins to IAV-WSN but overlapping T cell epitopes. This ensure that neutralising antibody to the WSN-IAV surface proteins does not prevent infection and T cell reactivation. In this model, T cell responses are associated with immune protection^21^. As expected, IAV-memory mice were protected from weight loss following infection with X31-IAV, Supplementary Figure 7A.

We analysed cytokine production by memory T cells before and 5 days after re-infection. The numbers of cytokine+ (IFNγ, IL-2, TNF) CD4 and CD8 T cells increased only slightly following re-infection, Fig 8A-B, Supplementary Figure 7B-C. Re-infection also led to an increase in the percentages of cytokine+ cells that were in cell cycle, most notably in cells in the spleen and dLN, Figure 8C-D, Supplementary Figure 7D-E. This was the case for both CD4 and CD8 T cells and regardless of which cytokine we examined; we did not analyse IL-2+ CD8 T cells as the numbers were too low. Most of the cytokine+ T cells were not, however, positive for Ki67, reflecting the low increase in cell number.

**Figure 8:**
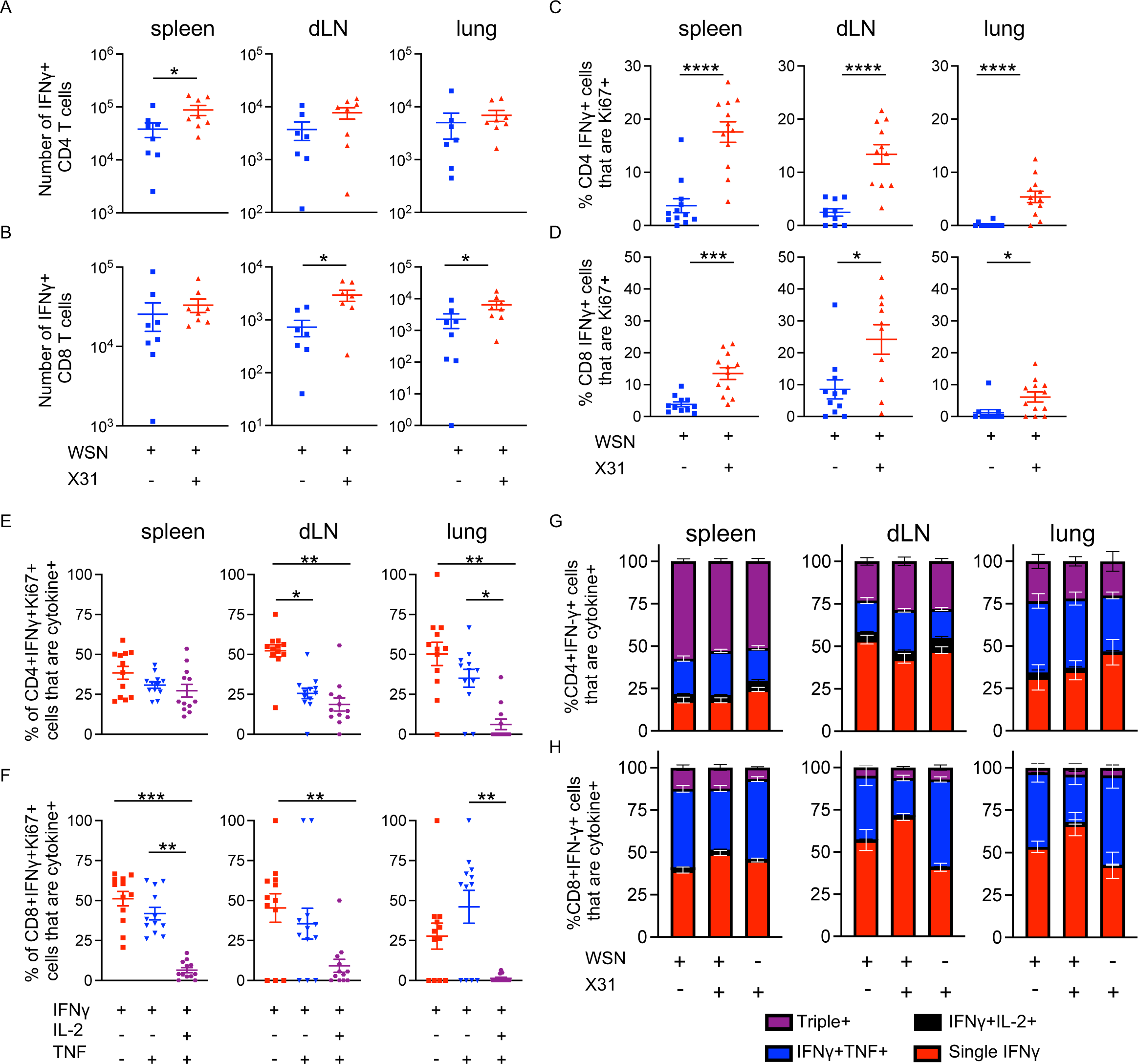
Triple cytokine+ T cells are less likely to be in cell cycle than single IFNγ+ T cells following challenge infection. C57BL/6 mice were infected i.n. with IAV on day 0. On day 30, some of these animals were infected with X31 IAV i.n. and 5 (A-F) or 35 days (G-H) later all mice were injected i.v. with fluorescently labelled anti-CD45 3 minutes prior to removal of organs. Single cell suspensions of spleens, mediastinal draining lymph node (dLN), and lung were activated for 6 hours in the presence of Golgi Plug by DCs incubated with IAV-Ag. CD45iv negative IAV specific IFNγ+ CD4 T cells (A,C,E,G) or CD8 T cells (B,D,F,H) were detected by flow to detect either IFNγ+ cells or IFNγ+ T cells that were Ki67+. Data are from two independent experiments with a total of 7-8 mice. Symbols represent each mouse and the horizontal line shows the mean of the group (A-F), in G-H error bars are SEM. In A-D significance tested by Mann-Whitney. In E-F, significance tested via ANOVA and Dunn’s. In all graphs: *: p<0.05, **:p<0.01, ***:p<0.001, ****:p<0.0001.

We next examined whether the IFNγ+ Ki67+ T cells were skewed towards single or multifunctional T cells, Figure 8E-F. For CD4 T cells in the dLN and lung, single IFNγ+ and double IFNγ+TNF+ cells had higher expression of Ki67 than triple cytokine+ cells. We saw a similar pattern in CD8 T cells in the spleen, dLN and lung with triple cytokine+ least likely to be Ki67+.

The low levels of Ki67 in triple cytokine+ cells may suggest that these cells would be less prevalent following reactivation and that single or double positive T cells may become a more prominent population. This was not the case. The percentages of IFNγ+ cells that were also expressing IL-2 and/or TNF were the same before and at day 5, Supplementary Figure 7F and day 35 following re-infection, Figure 8G-H. Moreover, mice given a primary infection with X31 had similar proportions of single, double and triple+ cells at day 35; too few cells were present in the primary infected mice at day 5 to analyse.

These data suggest that while CD4 triple cytokine+ T cells may be less likely to proliferate following reactivation, they are equally, or potentially better, equipped to survive into the secondary memory pool. In contrast, the single and double cytokine+ T cells that do proliferate following re-infection may be more likely to undergo apoptosis. Together these data suggest that these cytokine+ cell populations have distinct roles to perform during secondary response.

## Discussion

Here we reveal extensive heterogeneity within the memory CD4 T cell pool in lymphoid organs and the lung. Transcriptomic heterogeneity was clearly present within cells with the capacity to produce IFNγ, the key cytokine most associated with protection from IAV^2, 7–9, 16–19^. Importantly, our data suggest that heterogeneity within the memory pool also covers a spectrum of cells that do not fit into our current memory T cell paradigms that were first defined based on cell migration patterns rather than functional capacity.

Using various approaches to track IAV specific CD4 T cells we have made a number of key novel findings. First, our data show that the capacity to produce cytokines marks T cells with an enhanced ability to survive into the memory pool. Identification of the factors and cell types that promote cytokine+ cells could define how we could increase the size of the memory T cell pool following vaccination.

Cytokine+ CD4 T cells in secondary lymphoid organs expressed higher levels of the IL-7 receptor, CD127, and these cells and cytokine+ cells in the lung expressed higher levels of the anti-apoptosis molecule, Bcl2 than cytokine negative cells. Moreover, cells that expressed multiple cytokines had the highest expression of these molecules and expressed a number of pro-survival molecules at mRNA level. IL-7, which promotes Bcl2 expression, is a well-established survival signal for memory CD4 T cells^44, 61^. Which signals promote expression of CD127 on the cytokine+ cells will be a key question to address in future studies.

Second, within the cytokine+ fraction of the memory pool, we consistently identified three populations based on the capacity to produce one or more of IFNγ+, IL-2 and TNF. These three populations expressed distinct chemokine receptors, different levels and types of markers of T cell activation, and pro-survival molecules. These data suggest that these cells may have defined anatomical niches perhaps as a driver or result of their differentiation. These differences may impact on the cells’ survival and, importantly, their responses to reinfection. Alternatively, the populations could be distinct temporal states with cells oscillating between these clusters depending on recent interactions with other cell types and/or soluble molecules within their environment. Regardless, the data indicate a breadth in phenotype and function that has important implications for discussions on how we can most usefully define memory CD4 T cells subsets to advance an integrated understanding of these cells^62^.

A caveat of our model is that detection of the cytokine producing cells involves a short *ex vivo* reactivation stage. We therefore cannot know whether the expression of these molecules is altered during the reactivation. Our previous data did, however, demonstrate that single, double, and triple cytokine+ populations could be identified as early as 2 hours after re-activation, suggesting that these three populations are reflective of *in vivo* CD4 T cells with distinct phenotypes and functions^36^.

Double IFNγ+/TNF+ CD4 memory T cells often displayed an intermediate phenotype between triple cytokine+ cells and single IFNγ+ cells, for example in expression of Ki67, PD1, ICOS, Myc and the cytokines themselves. We think it is an over-simplification to consider these cells an intermediatory phase between single and triple cytokine+ cells as the double cytokine+ cells had a distinct transcriptional signature. Notably, the transcriptional signature of the double IFNγ+/TNF+ has some overlap with Tfh cells and the recently described T resident helper cells, including expression of *Cxcr5* and *Bcl6*^39, 51^. Resident Tfh and memory-Tfh like cells have been shown to differentiate into multiple different effector subsets upon re-activation *in vivo* suggesting these cells retain a level of plasticity^27, 51^. To test definitively the relationships between the three cytokine populations, comprehensive longitudinal lineage tracing approaches would be required.

We were surprised that the double and triple cytokine+ CD4 memory T cells did not dominate the secondary response to IAV. These cells expressed high level of transcripts for *Myc*, *Slc7a5* and the transferrin receptor, suggesting that they are poised to respond and proliferate following TCR activation^52^. We cannot exclude that memory triple or double cytokine+ CD4 T cells differentiate into single IFNγ+ during the re-infection. We think this is unlikely given the small changes in cell number and the stability of proportions of triple/double/single cytokine+ cells at days 5 and 30 following IAV re-infection. While there has been limited analysis of tertiary memory CD4 T cells, Bresser *et al* recently demonstrated that central memory CD8 T cell that have previously undergone the fewest rounds of division and have low expression of ‘effector cell’ genes dominate the secondary response^63^. In contrast, Wirth *et al* found that CD8 T cells can show increasing complexity following subsequent activation suggesting a contribution from multiple memory population to the tertiary pool^64^.

We identified IAV specific CD4 T cells using methods that restrict the population to a single immunodominant epitope or to T cells responding to numerous IAV epitopes. Our results are consistent between these methods suggesting that TCR specificity does not influence T cell survival into the memory pool. This conclusion is reinforced by finding the same TCR clones in several of the clusters within the single-cell RNAseq dataset. We suggest that environment is more likely, therefore, to dictate cell fate, than signals through the TCR.

We have analysed IAV specific CD4 and CD8 T cells in the same animals highlighting some key differences between these cell types. Most notably, cytokine+ CD8 T cells display a very limited decline in lymphoid organs. This reflects similar findings in LCMV infected mice in which the numbers of CD8 memory T cells able to produce cytokine following peptide stimulation remained stable for over 900 days while memory CD4 T cells declined ^65^. A second distinction in our study was revealed by examining sensitivity to the immunodominant peptides. While CD8 T cells produced a maximal cytokine response regardless of peptide dose, CD4 T cells produced less cytokine at lower peptide doses. It will be important to extend these data, examining T cells responding to other epitopes.

We did find consistent patterns of proliferation in IAV specific CD4 and CD8 T cells. Single IFNγ+ CD4 and CD8 T cells were more likely to be Ki67+ at primary, memory and recall timepoints compared to triple or double cytokine+ T cells. These data suggest that the link between proliferation capacity and multi-functionality is common to both CD4 and CD8 T cells. Understanding the molecular mechanisms that underlie this link could, therefore, reveal pathways that can be manipulated to enhance both the CD4 and CD8 T cell responses in the context of infectious disease and cancer, or limit such responses in autoimmune pathologies.

## Materials and Methods

### Animals and Study design

The aim of this study was to understand how cytokine production by memory CD4 and CD8 T cells changes over time and following a challenge re-infection. We used an influenza virus infection model in wildtype and reporter mice and tracked responding cells using MHC tetramers, cytokine assays, and the reporter system in the TRACE transgenic mice^43^. A description of the experimental parameters, samples sizes, any samples that were excluded, and the statistical analysis are described in each figure legend. No specified randomisation was conducted. Analysis of data was conducted in an unbiased manner.

10 week old female C57BL/6 mice were purchased from Envigo (UK). TRACE male and female and female C57BL/6 mice were maintained at the University of Glasgow under specific pathogen free conditions in accordance with UK home office regulations (Project Licenses P2F28B003 and PP1902420) and as approved by the local ethics committee. TRACE mice have been described previously^43^.

### Infections

IAV was prepared and titered in MDCK cells. 10-14 week old female C57BL/6 and male and female TRACE mice were briefly anesthetised using inhaled isoflurane and infected with 150-200 plaque forming units of IAV strain WSN in 20μl of PBS intranasally (i.n.). These mice are between 18-25g at the start of the experiment. Infected mice were rechallenged with 200PFU of X31. Infected mice were weighed daily for 14 days post-infection. Any animals that lost more than 20% of their starting weight were humanely euthanised. TRACE mice were given Dox+ chow (Envigo) for a total of 12 days starting two days prior to infection.

### Tissue preparation

Mice were injected intravenously (i.v.) with 1μg anti-CD45 (30F11, either labelled with Alexa 488 or PE (ThermoFisher, RRID: AB_2848416, RRID: AB_465668) 3 minutes before being euthanized by cervical dislocation. Spleen and mediastinal lymph nodes were processed by mechanical disruption. Single cell suspensions of lungs were prepared by digestion with 1mg/ml collagenase D (Sigma) and 30μg/ml DNAse (Sigma) for 40 minutes at 37°C in a shaking incubator followed by mechanical disruption. Red blood cells were lysed from spleen and lungs using lysis buffer (ThermoFisher).

### Ex vivo reactivation for cytokine

Bone marrow DCs were prepared as described^66^. Briefly, bone marrow cells were flushed from the tibias and femurs of female C57BL/6 mice and red blood cells removed. Cells were cultured in complete RPMI (RPMI with 10% foetal calf serum, 100μg/ml penicillin-streptomycin and 2mM L-glutamine) at 37°C 5% CO_2_ in the presence of GM-CSF (prepared from X-63 supernatant^67^) with media supplemented on day 2 and replaced on day 5. On day 7, DCs were harvested, incubated overnight with IAV antigen (MOI of 0.3) prepared as described^36^. Alternatively, DCs were incubated with NP peptides for 2 hours prior to co-cultures. Single cell suspensions of *ex vivo* organs were co-cultured with DCs in complete RMPI at a ratio of approximately 10 T cells to 1 DC in the presence of Golgi Plug (BD Bioscience). Co-cultures were incubated at 37°C, 5% CO_2_ for 6 hours.

### Flow cytometry staining

Single cell suspension were stained with PE or APC-labelled IA^b^/NP_311-325_ or APC labelled D^b^/ NP_368-374_ tetramers (both from NIH tetramer core) at 37°C, 5% CO_2_ for 2 hours in complete RPMI (RPMI with 10% foetal calf serum, 100μg/ml penicillin-streptomycin and 2mM L-glutamine) containing Fc block (24G2). Anti-CX3CR1 BV711 (SA011F11, BioLegend, RRID: AB_2565939) was added with the MHC tetramers. Surface antibodies were added and the cells incubated for a further 20minutes at 4°C. Antibodies used were: anti-CD4 APC-Alexa647 (RM4-5, ThermoFisher RRID: AB_1272183), anti-CD8 BUV805 (53-6.7, BD Bioscience RRID: AB_2870186), anti-CD44 BUV395 (IM7, BD Bioscience RRID: AB_2739963), anti-PD-1 PeCy7 (29F.1A12, BioLegend, RRID: AB_10696422), anti-PD1 BV605 (29F.1A12, BioLegend, RRID: AB_11125371), anti-ICOS PerCP-Cy5.5 (7E.17G9, BioLegend, RRID: AB_2832418), anti-CD127 APC (A7R34, ThermoFisher, RRID: AB_469435), CD103 PeCy7 (2E7, BioLegend, RRID: AB_2563690), anti-CD69 PerCP-Cy5.5 (H1.2F3, ThermoFisher, RRID: AB_1210703) and ‘dump’ antibodies: B220 (RA3-6B2, ThermoFisher, RRID: AB_1548761), F4/80 (BM8, ThermoFisher, RRID: AB_1548747) and MHC II (M5114, ThermoFisher, RRID: AB_1272204) all on eFluor-450. Cells were stained with a fixable viability dye eFluor 506 (ThermoFisher). For normalisation of CD127 and Bcl2 expression, the MFI of the EYFP+ cells was divided by the MFI of the naïve (CD44lo) CD4 T cells in the same animal in the same organ.

For intracellular staining, cells were fixed with cytofix/cytoperm (BD Bioscience, RRID: AB_2869013) for 20 minutes at 4°C and stained in permwash buffer with anti-cytokine antibodies for one hour at room temperature (anti-IFN-γ PE (XMG1.2, ThermoFisher RRID: AB_466193) or Brilliant violet 785 (XMG1.2, BioLegend, RRID: AB_11219004), anti-TNF Alexa-Fluor-488 (MP6-XT22, ThermoFisher, RRID: AB_469936) or Brilliant Violet 605 (MP6-XT22, BioLegend, RRID: AB_11123912), anti-IL-2 APC (JES6-5H4, ThermoFisher, RRID: AB_469490) or Brilliant violet 711 (JES6-5H4, BioLegend, RRID: AB_2564225), Bcl2 PeCy7 (Blc/10C4, BioLegend, RRID; AB_2565246), anti-Ki67 PeCy7 (16A8, BioLegend,) and cells washed with permwash buffer. To detect Myc, cells were first stained with unlabelled anti-Myc (D84C12, Cell Signalling Technologies, RRID: AB_1903938), cells washed with permwash and then stained with PE-anti-rabbit (Cell signalling Technologies, RRID:AB_2799931). Stained cells were acquired on a BD LSR or Fortessa and analysed using FlowJo.

### BD Rhapsody single cell RNA-seq

CD4 T cells from spleens and lungs of IAV infected TRACE were isolated by CD4 negative selection following the manufacturer’s instructions (StemCell, CD4 Isolation Kit). The cells were activated with IAV-Ag-DCs for 4 hours and then stained with anti-CD4 APC-Alexa647, anti-CD44-PerCP-Cy5.5 (IM7, ThermoFisher, RRID:AB_925746), anti-MHCII-e450, anti-B220-e450, CD8-e450 (53-6.7, ThermoFisher, RRID:AB_1272198), F4/80-e450, and CD45 Sample Tags to enable multiplexing (BD Bioscience, RRID: AB_2870301) and AbSeq antibodies to ICOS (AMM2072, RRID: AB_2876063) and PD1 (AMM2138, RRID: AB_2876215). CD44^hi^/EYFP+ and CD44^lo^/EYFP negative cells from combined spleen or lung samples were FACS sorted on an ARIA IIU and cells transferred into BD Rhapsody Sample Buffer and loaded onto a scRNA-seq Rhapsody Cartridge (5000 EYFPnegative CD44^lo^ spleen cells; 5000 EYFP+CD44^hi^ spleen cells; and 1000 EYFP+CD44^hi^ lung cells. Manufacturer’s instructions (Mouse VDJ CDR3 library preparation) were followed to prepare libraries for sequencing with the BD Rhapsody Immune response targeted panel (mouse, RRID: AB_2870301), additional custom made primers (Supplementary excel file 1), sample Tags and VDJ CDR3. Pair-end sequencing was performed by Novogene on an Illumina MiSeq PE250.

### Single cell RNA-seq analysis

Data were initially processed on SevenBridges and then analysed predominantly using Seurat (V4.2.0)^68^ and scRepertoire(V1.7.2)^69^. Briefly, to perform dimensionality reduction: count data was normalised using scTransform prior to principal component analysis, Uniform Manifold Approximation and Projection (UMAP) dimensional reduction, Nearest-Neighbour graph construction and cluster determination within the Seurat package, and dimensionality reduction performed. Log normalising and scaling of the count data was conducted prior to differential gene expression analysis. To test for differential gene expression, the FindAllMarkers function was used (min.pct = 0.4 & minLogFC = 0.25, model.use = MAST). Only genes with a Bonferroni corrected p value <0.05 were considered statistically different. For TCR analysis, clonal expansion was calculated based on the same TCR being detected once (Single Clone) or multiple times within the same animal. All TCR analysis utilised stock or customised code from the scRepertoire package. Further packages used for data analysis, organisation and visualisation included: workflowR^70^, dplyr, ggplot2, cowplot, ggVenn, circlize^71^, plot1cell^72^ and ComplexHeatmap^73^. All code used can be found at https://github.com/JonathanNoonan/Westerhof_2023

### Statistical analysis

All data other than scRNAseq were analysed using Prism version 9 software (GraphPad). Differences between groups were analysed by unpaired ANOVAs, T-tests or Mann-Whitney test as indicated in figure legends. Multi-group non-parametric data were analysed using Kruskal-Wallis testing with a *post hoc* Dunn’s test. In all figures * represents a p value of <0.05; **: p<0.01, ***: p<0.001, ****: p<0.0001.

## Data availability

scRNA-seq data have been deposited at GEO and are publicly available as of the date of publication; GSE220588.

## Acknowledgments

We thank the staff within the School of Infection and Immunity Flow Cytometry Facility and Biological Services at the University of Glasgow for technical assistance. Thank you to members of the Kurowska-Stolarska for technical assistance with the BD Rhapsody experiment. Thank you to Prof James Brewer and Dr Edward Roberts for critical review of the manuscript and the MacLeod lab for discussions. We thank the NIH tetramer core facility for the provision of IA^b^-NP_311-325_ and D^b^/NP NP_368-74_ tetramers. The work was funded by a GLAZgo Discovery Centre PhD fellowship to LMW, a Marie Curie Fellowship (334430) and a Wellcome Trust Investigator Award (210703/Z/18/Z) to MKLM.

## CRediT authorship contribution statement

LMW: conception, investigation, formal analysis, visualization, writing - original draft and reviewing/editing. JN: formal analysis, data curation, software, visualization, writing - reviewing and editing. KEH: investigation, formal analysis, writing - reviewing and editing. ETC: investigation, formal analysis, writing - reviewing and editing, ZC: investigation, formal analysis, writing-reviewing and editing, TP: investigation, writing - reviewing and editing. MRJ: formal analysis, writing – reviewing and editing. NB: software, writing – reviewing and editing. MKLM: Supervision, project management, funding acquisition, conception, investigation, formal analysis, visualization, writing-original draft and reviewing.

## Supplemental tables

**Supplementary Table 1**: Differentially expressed genes from the scRNAseq analysis Supplementary File 1: Sample Tag sequences, TCR primers and additional genes for the scRNAseq analysis

