## Supplementary Figures for "Enhanced survival and low proliferation marks multifunctional virus-specific memory CD4 T cells"

Supplementary Figure 1: Gating strategies for identification of IAV specific CD4 and CD8 T cells and example staining for surface markers on lung T cells

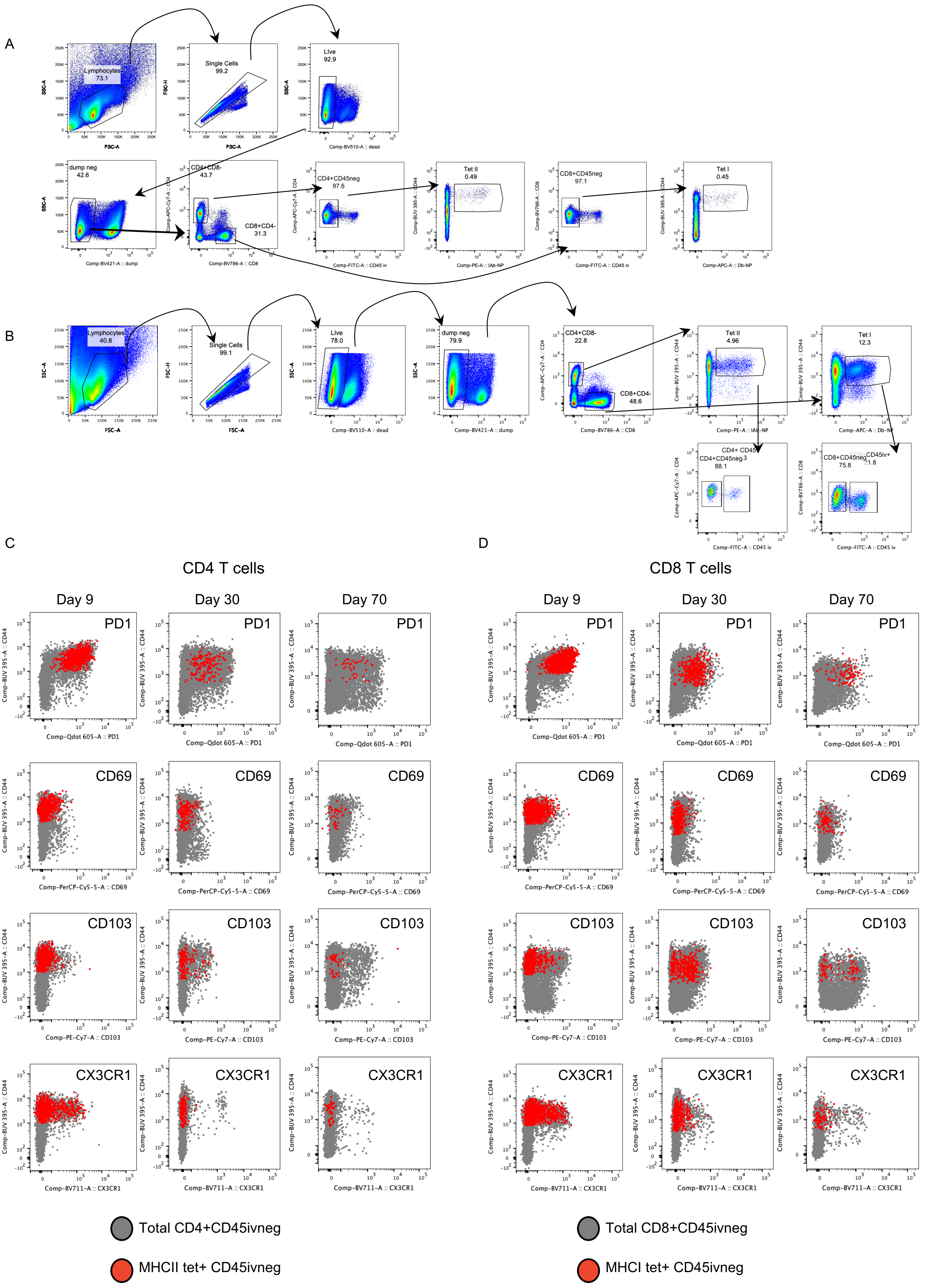

**Supplementary Figure 1**

*Example gating of IAV specific CD4 and CD8 T cells identified by MHC tetramers*

C57BL/6 mice were infected i.n. with IAV on day 0 and injected i.v. with fluorescently labelled anti-CD45 3 minutes prior to removal of organs for analysis. Single cell suspensions of spleen (A) and lung (B) are shown at day 9 of infection. A, shows example gating used in most figures in which we gate on CD45 i.v. negative tetramer+ or cytokine+ T cells. Example FACS plots of splenic CD45iv negative MHC tetramer+ cells CD4 (C) and CD8 (D) (shown in red) in comparison to total CD45iv negative T cells (grey).

Supplementary Figure 2: IL-2 and TNF+ CD4 and CD8 T cells show minimal decline between day 9 and day 30 in secondary lymphoid organs

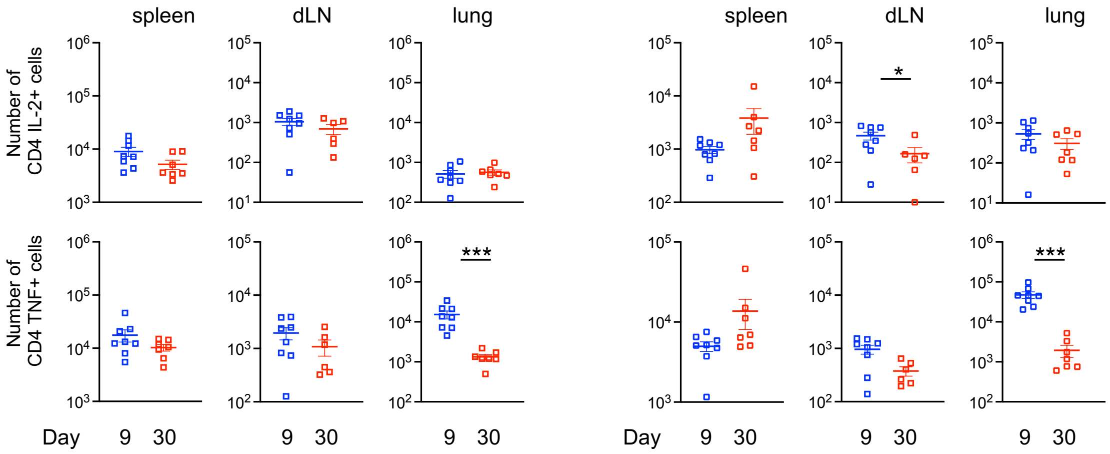


**Supplementary Figure 2**

IL-2 and TNF+ CD4 and CD8 T cells show minimal decline between day 9 and day 30 in secondary lymphoid organs

C57BL/6 mice were infected i.n. with IAV on day 0 and injected i.v. with fluorescently labelled anti-CD45 (CD45iv) 3 minutes prior to removal of organs. Single cell suspensions of spleens, mediastinal draining lymph node (dLN), and lung were examined after 9 or 30 days and either stained with MHCII/NP or MHCI/NP tetramers, or activated with IAV-peptide loaded DC. Numbers of IL-2 or TNF+ CD4 (A) or CD8 T cells (B) were calculated. Data are from two independent time course experiments with a total of 7-8 mice/time point. Y-axis set at the limit of detection and errors are SEM. Significance tested by a Mann-Whitney, *: $p<0.05$, ***:$p<0.001$.

Supplementary Figure 3: TRACE mice enable identification of CD4 T cells responding to IAV infection

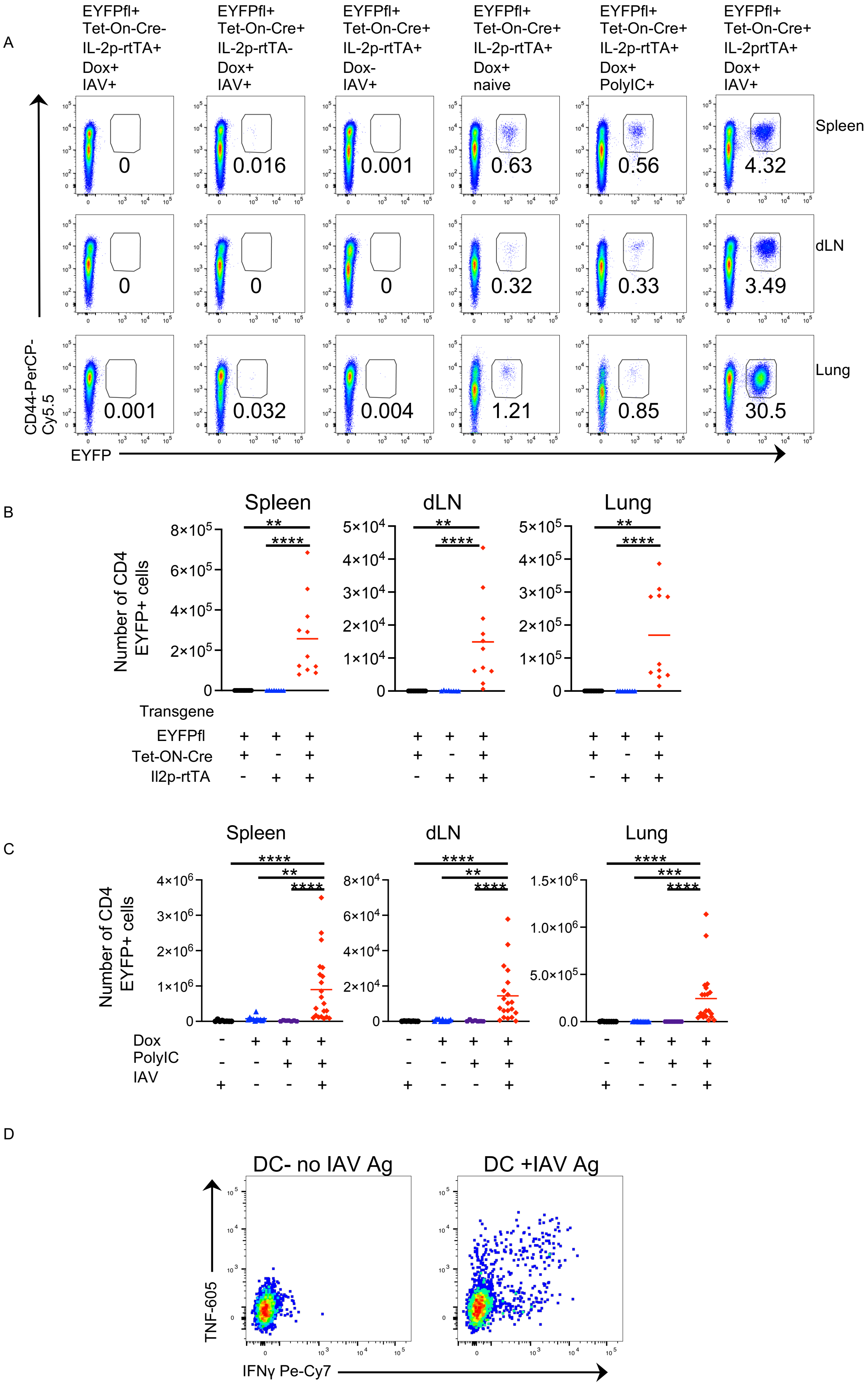

**Supplementary Figure 3**
*TRACE mice enable identification of CD4 T cells responding to IAV infection*

TRACE mice that are positive for all three transgenes and double transgenic mice expressing STOP floxed EYFP (EYFPfl) and either tet-ON-Cre or the IL-2 promoter driving rtTA (IL-2p-rtTA) were given doxycycline (Dox) diet for a total of ten days starting two days prior to intranasal infection with IAV or instillation of 20μg of PolyIC intranasally as indicated. In A, cells are gated as in Supplementary Figure 1A on CD4+ live lymphocytes that are negative for CD8, B220, MHCII and F4/80 in the indicated organ 8 days after infection/polyIC treatment and the numbers show the percentages of EYFP+ cells within the gate. In B, mice with the indicated transgenes were given dox diet and the numbers of EYFP+ CD4 T cells examined 8-12 days following treatment, data are combined from 3 independent experiments with 2-5 mice per experiment. In C, TRACE mice were treated/infected as indicated and the numbers of EYFP+ CD4 T cells examined 8-12 days following infection, data are combined from 4 independent experiments with 2-10 mice per experiment. Significance tested by a Kruskal-Wallis test followed by a Dunn's multiple comparison test: *: $p<0.05$, **:$p<0.01$, ***:$p<0.001$, ****:$p<0.0001$. D shows example staining from a day 8 IAV infected mouse, the cells on the left were co-cultured with DCs that had not received IAV-Ag, the plot on the right shows cells from the same mouse co-cultured with IAV-Ag+ DCs for 6 hours. Cells are gated on live single CD4+ EYFP+ cells that are CD45iv negative.

Supplementary Figure 4: scRNAseq reveals heterogeneity in the memory CD4 T cell pool

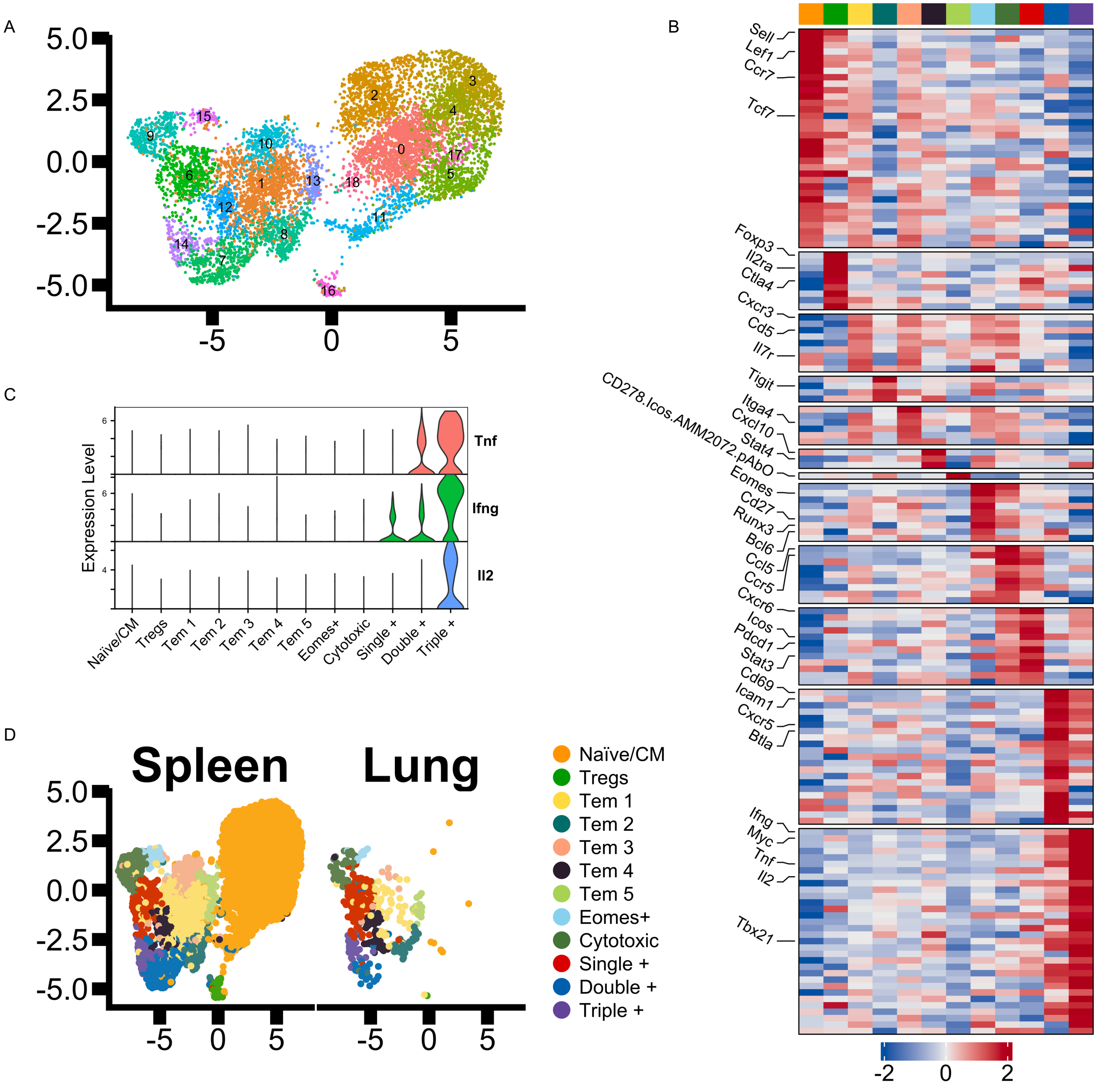


**Supplementary Figure 4**

*scRNAseq reveals heterogeneity in the memory CD4 T cell pool*

TRACE mice were infected with IAV and injected with anti-CD45 3 minutes prior to removal of spleens and lungs. Isolated CD4 T cells were activated for 4 hours by co-culture with IAV-Ag DCs and the CD45iv negative CD4+CD44$^{hi}$EYFP+ cells (spleens and lung) and CD4+CD44$^{lo}$EYFPnegative (spleens) cells were FACS sorted and their transcriptomes examined by scRNAseq. UMAP of eight naïve/central memory clusters, one Treg cluster, and 10 memory clusters (A). DEGs in main clusters identified by comparison between the indicated populations and all other clusters (B). Where genes were differentially expressed in multiple clusters, the gene was visualised in the cluster with the highest fold change; for a list of all DEGs see Supplementary Table 1. Expression of *Tnf*, *Ifng* and *Il2* by each cluster (C). UMAP of analysed cells displaying cells from the spleen and lung separately (D).

Supplementary Figure 5: Triple cytokine+ T cells make more cytokine on a per cell basis than single cytokine+ T cells

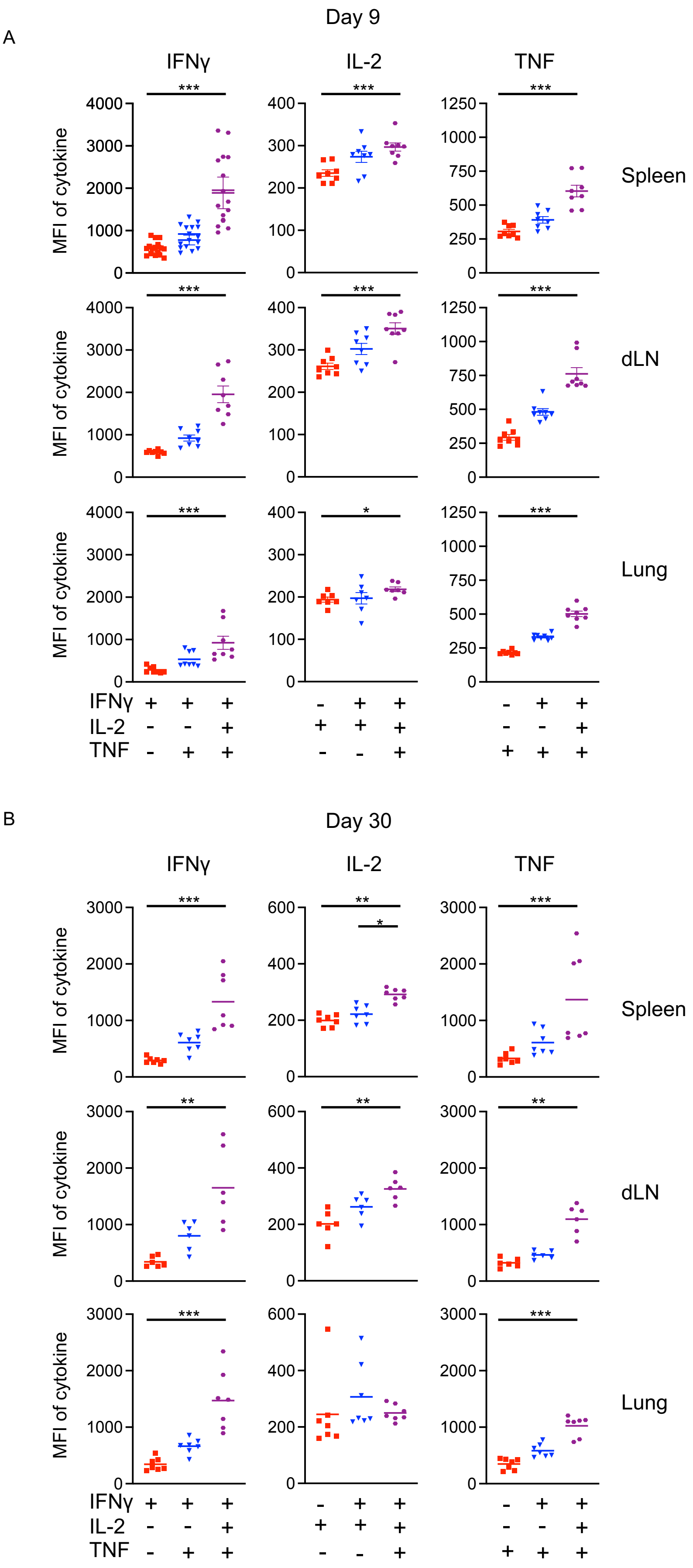

**Supplementary Figure 5**

*Triple cytokine+ CD4 T cells produce more cytokine on a per cell basis than single cytokine+ T cells*

C57BL/6 mice were infected i.n. with IAV on day 9 or day 30 and injected i.v. with fluorescently labelled anti-CD45 3 minutes prior to removal of organs for analysis. Single cell suspensions of spleens, mediastinal draining lymph node (dLN), and lung were activated by bmDCs incubated with IAV-Ag. CD45iv negative IAV specific CD4 T cells cytokine+ at day 9 (A) or day 30 (B) T cells were detected by flow cytometry. Each symbol represents a mouse and the line shows the mean of the group with SEM shown. Significant differences were assessed by Friedman's multiple comparison test followed by multiple comparisons with Dunn's multiple comparison test; *: $p<0.05$, **:$p<0.01$, ***:$p<0.001$.

Supplementary Figure 6: Single IFNγ+ CD4 memory T cells express higher levels of PD1 and ICOS than triple cytokine+ cells

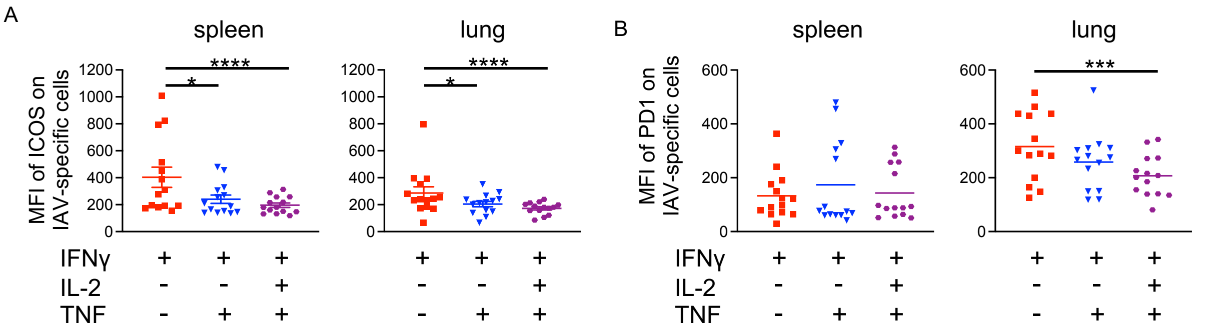


**Supplementary Figure 6**
Single IFNγ+ CD4 memory T cells express higher levels of PD1 and ICOS than triple cytokine+ cells

C57BL/6 mice were infected i.n. with IAV on day 0 and injected i.v. with fluorescently labelled anti-CD45 3 minutes prior to removal of organs at day 40. Single cell suspensions of spleens and lung were activated by DCs incubated with IAV-Ag preparation. CD45iv negative cytokine+ CD4 T cells were detected by flow cytometry to detect PD1 (A) and ICOS (B) expression. Data are from two separate experiments. Each symbol represents a mouse and the horizontal line shows the mean of the group. Significance tested via paired Friedman analysis with Dunn's multiple comparison test *: p<0.05, ***:p<0.001, ****:p<0.0001.

Supplementary Figure 7: Previous infection with IAV leads to a protective response following re-challenge infection but a limited increase in the number of cytokine+ T cells

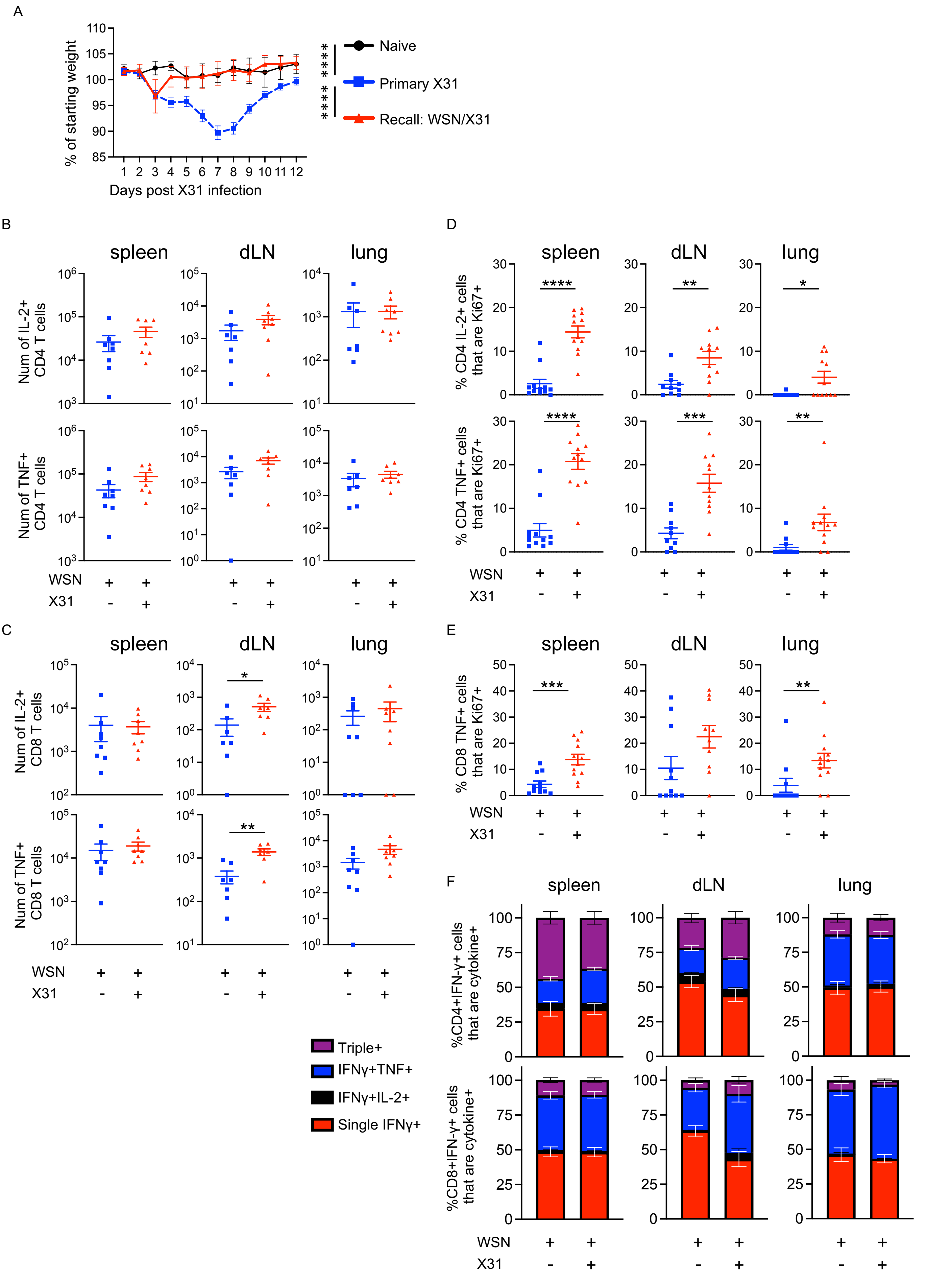

**Supplementary Figure 7**

*Previous infection with IAV leads to a protective response following re-challenge infection but a limited increase in the number of cytokine+ T cells*

C57BL/6 mice were infected i.n. with WSN IAV on day -30 and then infected with X31 on day 0. Controls were age-matched naïve animals or naïve mice infected with X31 IAV on day 0. Mice were weighed and the difference in weight loss calculated by measuring the area and the curve (A). On day 5 after the challenge infection, a separate cohort of mice were injected i.v. with fluorescently labelled anti-CD45 3 minutes prior to removal of organs for analysis. Single cell suspensions of spleens, mediastinal draining lymph node (dLN), and lung were activated by bmDCs incubated with IAV-Ag preparation. CD45iv negative IAV specific IL-2+ of TNF+ CD4 T cells (B) or CD8 T cells (C) were detected by flow cytometry and their expression of Ki67 examined (D,E). The proportions of IFN$\gamma$+ CD4 and CD8 T cells expressing IL-2 and TNF were calculated (F). In A, naïve animals are from one experiment (4 naïve animals) and infected animals combined from two experiments with a total of 12 primary infected animals and 9 re-infected animals. Symbols are the means of the groups and error bars show SEM. The areas under the curves were compared by ANOVA followed by a Tukey's multiple comparison test with ****:sp<0.0001. In B-E, symbols represent each mouse and the horizontal line shows the mean of the group. Significance tested by Mann-Whitney, *: p<0.05, **:p<0.01, ***:p<0.001. In F, error bars are SEM.
